# Acid stress signals are integrated into the σ^B^-dependent general stress response pathway via the stressosome in the food-borne pathogen *Listeria monocytogenes*

**DOI:** 10.1101/2021.12.20.473419

**Authors:** Duarte N. Guerreiro, M. Graciela Pucciarelli, Teresa Tiensuu, Diana Gudynaite, Aoife Boyd, Jörgen Johansson, Francisco García-del Portillo, Conor P. O’Byrne

## Abstract

The general stress response (GSR) in *Listeria monocytogenes* plays a critical role in the survival of this pathogen in the host gastrointestinal tract. The GSR is regulated by the alternative sigma factor B (σ^B^), whose role in protection against acid stress is well established. However, the mechanisms leading to its activation by low pH are unknown. Here, we investigated the involvement of the stressosome, a sensory organelle, in transducing low pH signals to induce the GSR. Mild acid shock (15 min at pH 5.0) activated σ^B^ and conferred protection against a subsequent lethal pH challenge. A mutant strain where the stressosome subunit RsbR1 was present but its remaining paralogues were genetically inactivated retained the ability to induce σ^B^ activity at pH 5.0. The role of stressosome phosphorylation in signal transduction was investigated by mutating the putative phosphorylation sites in the core stressosome proteins RsbR1 (*rsbR1* T175A, T209A, T241A) and RsbS (*rsbS* S56A), or in the active site of the stressosome kinase RsbT (*rsbT* N49A). The *rsbS* S56A and *rsbT* N49A mutations abolished the response to low pH. The *rsbR1* T175A variant, retained a near-wild type phenotype. The *rsbR1* T209A and *rsbR1* T241A mutants displayed constitutive σ^B^ activity. Mild acid shock upregulates invasion genes and stimulates epithelial cell invasion, effects that were abolished in mutants with an inactive or overactive stressosome. Overall, the results show that the stressosome is required for acid-induced activation of σ^B^ in *L. monocytogenes*. Furthermore, RsbR1 can function independently of its paralogues and that signal transduction requires RsbT-mediated phosphorylation of RsbS on S56 and RsbR1 on T209. These insights shed light on the mechanisms of signal transduction that activate the GSR in *L. monocytogenes* in response to acidic environments, and highlight the role this sensory process in the early stages of the infectious cycle.

**Author summary:** The stress sensing organelle known as the stressosome, found in many bacterial and archaeal lineages, plays a crucial role in both stress tolerance and virulence in the food-borne pathogen *Listeria monocytogenes*. However, the mechanisms that lead to its activation and the subsequent activation of the general stress response have remained elusive. In this study, we examined the signal transduction mechanisms that operate in the stressosome in response to acid stress. We found that only one of the five putative sensory proteins present in *L. monocytogenes*, RsbR1, was required for effective transduction of acid tress signals. We further found that phosphorylation of RsbS and RsbR1, mediated by the RsbT kinase, is essential for signal transduction. Failure to phosphorylate RsbS on Serine 56 completely abolished acid sensing by the stressosome, which prevented the development of adaptive acid tolerance. The acid-induced activation of internalin gene expression was also abolished in mutants with defective stressosome signalling, suggesting a role for the stressosome in the invasion of host cells. Together the data provide new insights into the mechanisms that activate the stressosome in response to acid stress and highlight the role this sensory organelle plays in virulence.

## Introduction

The firmicute *Listeria monocytogenes* is a stress-tolerant Gram-positive bacterium that causes listeriosis, a food-borne infection that poses a particular risk for immunocompromised individuals and for pregnant women that is often associated with a high mortality rates (ranging from 20 to 30%) (1). The ability of *L. monocytogenes* to survive and proliferate in a broad range of harsh environments, including those found in food processing facilities and the human gastrointestinal (GI), tract makes it a particular problem for producers of ready-to-eat foods (2–4). Following the entry into the host, *L. monocytogenes* can withstand an array of potentially lethal conditions in the GI tract, such as the extreme acid pH of the stomach, high osmotic pressure and high concentration of bile salts in the duodenum (reviewed in (5, 6). *L. monocytogenes* employs the alternative sigma factor B (σ^B^) to upregulate a regulon of approximately 300 genes, which collectively confer an array of homeostatic and protective mechanisms that allow survival under severe stress conditions (reviewed in (7, 8). *L. monocytogenes* relies on *σ*^B^ to enhance survival in the GI tract and to establish infection (9). A Δ*sigB* mutant exhibits attenuated virulence (10–12). This has been partly attributed to the σ^B^-dependent expression of the internalins A and B, encoded by *inlA and inlB,* respectively. These surface-expressed invasins promote internalization of *L. monocytogenes* into epithelial cells expressing the E-cadherin and C-Met receptors, respectively (13–15). Interestingly, exposure of *L. monocytogenes* to acid shock increases *inlA* and *inlB* expression and invasiveness in epithelial cells (14,16,17). Additionally, loss of *sigB* is associated with a marked reduction in survival at extreme low pH (18). The σ^B^ regulon is upregulated by mild stress conditions (19–24). Although important for *L. monocytogenes* acid tolerance, Δ*sigB* mutants are still able to develop further resistance towards lethal acid pH when pre-exposed to mild acidic stress (pH 5.0) for one hour (25–27). This phenomenon is known as the adaptive acid tolerance response (ATR) (28, 29). While it is well established that σ^B^ plays a critical role in acid tolerance its role in the ATR is poorly understood. Genes known to be involved in acid tolerance are upregulated in a σ^B^-dependent manner following a mild acid treatment (23,30–32) but the mechanisms of signal transduction involved have not been described yet.

The regulatory pathway of σ^B^ activation in *L. monocytogenes* was initially inferred from *Bacillus subtilis* due to the high degree of conservation between the *sigB* operons of these two genera, composed of *rsbR, rsbS, rsbT, rsbU, rsbV, rsbW, sigB* and *rsbX* (33, 34). These operons encode components of a signal transduction cascade responsible for triggering the dissociation of σ^B^ from RsbW (anti-sigma factor) during the onset of stress, allowing the formation of the holoenzyme RNA polymerase:σ^B^ (*E*σ^B^) and consequently the upregulation of the σ^B^ regulon (35, 36) (Fig 1A). At the top of the signal transduction pathway is a high molecular weight protein complex (∼1.8 MDa) called the stressosome that serves as a sensory hub, integrating environmental signals into the σ^B^ pathway (Fig 1A) (37–39). Genes encoding this macromolecular complex are found in a phylogenetically diverse range of species in both bacteria and archaea (40). The stressosome functions as an environmental and nutritional stress sensor in *L. monocytogenes* (37,41–43). The *L. monocytogenes* stressosome is composed of RsbR1, an orthologue of *B. subtilis* RsbRA, and its paralogues RsbR2 (Lmo0161), RsbL (Lmo0799), RsbR3 (Lmo1642) and potentially RsbR4 (Lmo1842), although the latter has not yet formally been shown to be part of the stressosome complex. Interestingly, the N-terminal domains of the RsbR paralogues of *L. monocytogenes* and *B. subtilis* share little amino acid conservation, suggesting that the sensory roles of these proteins may differ between species. These proteins combine with the core protein RsbS and the serine-threonine kinase RsbT to form a pseudo-icosahedral complex with a 2:1:1 ratio (RsbR1:RsbS:RsbT) (44) (Fig 1A). The current model suggests that the N-terminal domains of RsbR1, which form protruding turrets, are responsible for sensing environmental stresses. Following signal detection it is thought that a structural rearrangement propagates into the stressosome core leading to the activation of kinase RsbT, which phosphorylates RsbR1 at T209 and RsbS at serine 56 (S56), promoting the release of RsbT from the stressosome core and signal cascade initiation (39, 44). Thus far, this model does not takes into account interactions among the RsbR1 paralogues, although a recent study suggests that they may act negatively on the stressosome assembly (42). It is currently unknown if RsbR1 and its paralogues are able to form heterodimers within the stressosome, although in *B. subtilis* YtvA dimers (the homologue of RsbL) can form heterotetramers with RsbRA dimers (45).

**Fig 1.**
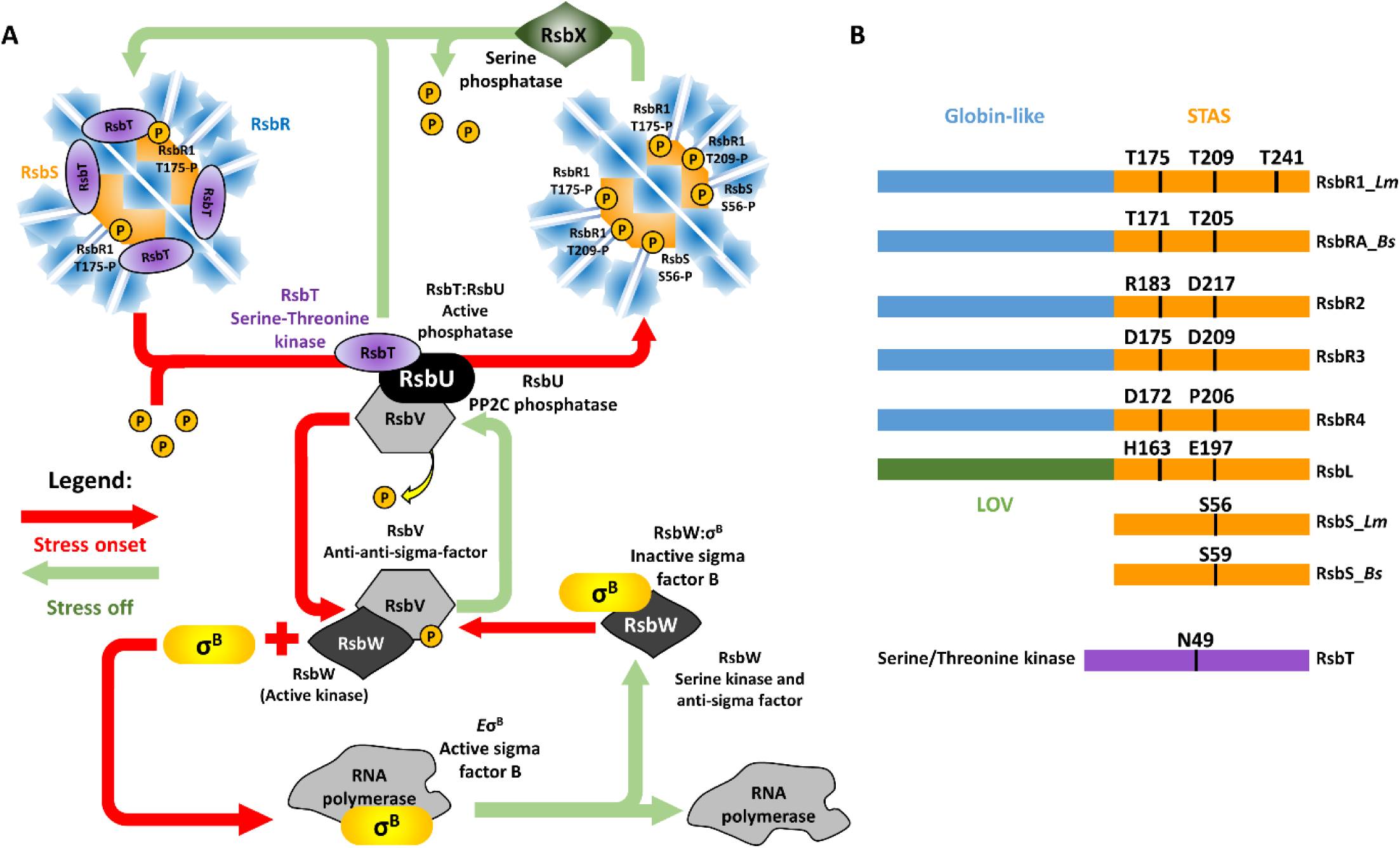
Model depicting the stressosome-mediated regulation of σ^B^ and alignment of stressosome proteins. (A) The putative sensory proteins RsbR1 and its paralogues RsbR2, RsbR3, RsbL, the scaffold protein RsbS and the serine-threonine kinase RsbT form the stressosome. Environmental/nutritional stress is hypothesized to be sensed by one or more RsbR1 paralogue via its N-terminal non-haem binding globin domains forming the turrets at the stressosome surface (blue). The stressosome core (orange) is composed of the STAS domains of both RsbS and RsbR1 C-termini. Stress triggers T209 phosphorylation in RsbR1 and S56 in RsbS, freeing RsbT (purple) from the stressosome core. The phosphatase RsbU interacts with the free RsbT, activating its phosphatase activity towards RsbV. Once dephosphorylated, the serine kinase and anti-sigma factor RsbW, which has higher affinity with the non-phosphorylated RsbV, releases σ^B^ and phosphorylates RsbV once more. σ^B^ is then able to interact with the RNApol forming the holoenzyme *E*σ*^B^* enabling the upregulation of the general stress regulon. The phosphatase RsbX is responsible for the deactivation of the stressosome by dephosphorylating its residues and enabling the re-sequestration of RsbT and the inactivation of the signal cascade. The model shown incorporates the current available data from studies on both *Listeria* and *Bacillus* (see text for citations). (B) Domain organization of *L. monocytogenes* RsbR1_*Lm* and its paralogues, RsbS_*Lm* and RsbT and *B. subtilis* RsbRA_*Bs* and RsbS_*Bs*. The N-terminal domains of RsbR1 and paralogues fold into non-haem binding globin-like domains that form the stressosome turrets, while the blue-light sensor RsbL folds into an LOV domain. The C-termini of RsbR1 and paralogues fold into STAS domains forming the stressosome core along with RsbS. The serine-threonine kinase RsbT responsible for the stressosome phosphorylation requires the ATP-Mg^2+^ chelating residue N49 for its kinase activity.

In the stressosome core the RsbR1/RsbRA C-terminal domain and RsbS, fold into <u>S</u>ulphate <u>T</u>ransporter and <u>A</u>nti-<u>S</u>igma factor antagonist (STAS) domains (46, 47). Several residues in the STAS domains of *B. subtilis* RsbRA (T171 and T205) and RsbS (S59) are phosphorylatable and display a critical role in the stressosome activation (48, 49). Similarly, *L. monocytogenes* RsbR1 also possesses putative phosphorylation sites at positions T175, T209 and T241, and S56 in RsbS (Fig 1B and 1C). However, these threonine residues are absent in the remaining paralogues of RsbR1 and so far, only RsbR1 T175 phosphorylation has been confirmed (42). The RsbT kinase is responsible for phosphorylation of all residues in the RsbR1 and RsbS and the inactivation or deletion of this protein results on the impairment of the σ^B^ activation in both *L. monocytogenes* and *B. subtilis* (41,42,50–52). Once the stress has dissipated or the bacterium has adequately responded to it, the stressosome is restored to the ground state by dephosphorylation of the core proteins, through the action of the serine phosphatase RsbX (50,53–56).

In this study, we examined the role of the *L. monocytogenes* stressosome in sensing mild acid stress and in activating the general stress response and conferring adaptive acid resistance. We constructed an array of mutations, where either the paralogues of RsbR1 were deleted, the kinase activity of RsbT was inactivated, or the putative phosphorylation sites in the stressosome core were substituted with alanine. Our results revealed that RsbR1 has the ability to activate σ^B^ in response to mild acid stress when the other four paralogues were absent. The phosphorylation sites in RsbR1 (T175 and T209) modulate the intensity of σ^B^ activity in the absence of stress, but have no role in the stressosome’s low pH sensing. Furthermore, the RsbS S56 phosphorylation site and the kinase activity of RsbT are crucial for σ^B^ activation at low pH. The extent of σ^B^ activation in response to mild acid stress correlates closely with the survival under lethal acidic conditions (pH 2.5). We also show that the stressosome is required for the induction of the internalins *inlA* and *inlB* under conditions of low pH stress and for the modulation of *L. monocytogenes* internalization into epithelial cells. Together, our results point to a central role for the stressosome in acid stress sensing, the development of acid resistance and in the regulation of host cell invasion, through the activation and modulation of σ^B^ activity. These findings point to the stressosome as a possible target for new antimicrobials to control this pathogen in food and in the gastrointestinal tract.

## Results

### The transcription of *lmo0596* is induced by acid in σ^B^- dependent manner

The σ^B^ regulon in *L. monocytogenes* is composed of a large number of genes that are heterogeneously expressed in response to the stress encountered (9). The expression of the gene *lmo2230*, which encodes a putative arsenate reductase, is well established as a reliable σ^B^-reporter (24,42,57), however we sought to use an additional σ^B^- reporter gene in this study. The gene *lmo0596* was a potential target, as it possesses a σ^B^ promoter and had been shown in other studies to belong to the σ^B^ regulon (9,58,59). Although, Lmo0596 has no predicted function, it is homologous to *Escherichia coli* HdeD, a transmembrane protein that plays a role in acid tolerance (60–63). *In vitro* transcription experiments showed that *lmo0596* transcription is σ^B^- dependent and PrfA-independent in *L. monocytogenes* (64), while *in vivo* studies showed it to be highly σ^B^-dependent (57, 58). In addition, acid treatment at pH 5.0 upregulates *lmo0596* within the first 15 min post stress (17). To validate *lmo0596* as a σ^B^-reporter, the *L. monocytogenes* wild type strain EGD-e and an isogenic Δ*sigB* mutant strain were grown to mid log-phase (OD_600_ = 0.4) (Fig 2A), and the transcription of *lmo0596* and *lmo2230* were then measured following acid treatment (pH 5.0). The transcript levels of both *lmo0596* and *lmo2230* increased after the acid treatment in the wild type strain, plateauing at 15 min, while transcript levels remained unchanged in the Δ*sigB* strain (Fig 2B and 2C and S1A Fig). The expression of *lmo0596* also increases as the pH decreases, reaching its maximum at pH ∼5.0-5.5 (S1B Fig). The presence of a putative PrfA box in the regulatory region upstream from *lmo0596* prompted us to examine whether PrfA might also influence its transcription. Northern blots used to measure *lmo0596* transcription in wild type and Δ*prfA* backgrounds showed that PrfA does not influence *lmo0596* expression under the conditions tested (S1A Fig). Overall, these results showed that *lmo0596* transcription was induced by mild acid treatment in a σ^B^-dependent manner, making it suitable to use as a reporter of σ^B^ activity.

**Fig 2.**
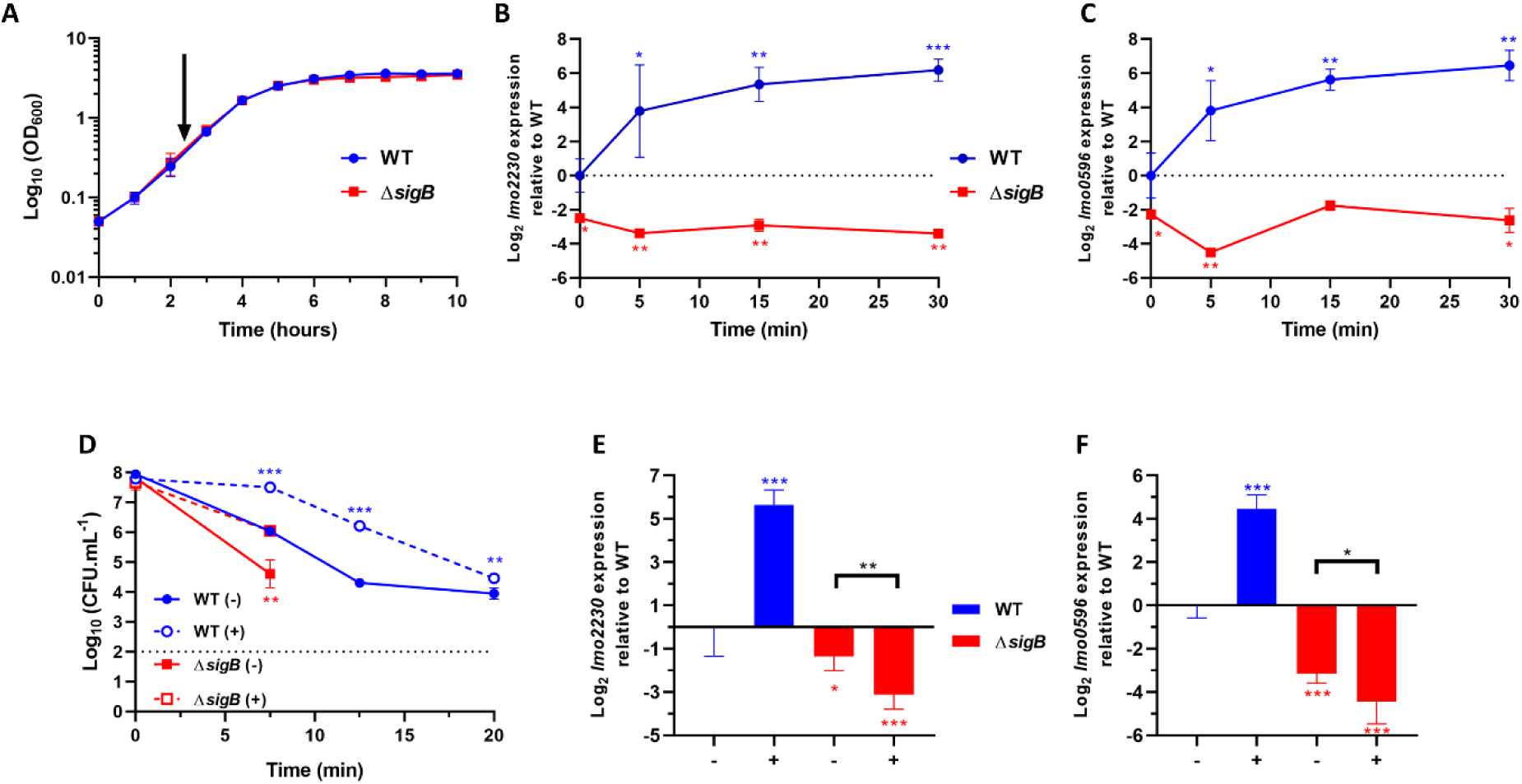
σ^B^-dependent transcription of *lmo2230* and *lmo0596* is upregulated rapidly under mild acid stress conditions. (A) Growth curves obtained from Log_10_ (OD_600_) measurements of wild type and Δ*sigB* strains in BHI at 37°C. The arrow points to OD_600_ = 0.4, the mid-log phase of *L. monocytogenes* in BHI. Time-lapse expression measurements of the genes (B) *lmo2230* and (C) *lmo0596* in wild type and Δ*sigB* strains grown to OD_600_ = 0.4 and subsequently treated at a mild pH 5.0 (HCl 5 M) at 0 min. Samples were taken at 0, 5, 15 and 30 min post treatment. (D) Mid-log phase cultures untreated at pH 7.0 (-) or treat at pH 5.0 (+) for 15 min and subsequently challenged in acidified BHI (pH 2.5). The dashed line represents the detection threshold. Samples were taken at 0, 7.5, 12.5 and 20 min. Survival data is expressed as Log_10_ (CFU.mL^-1^). Gene expression of *lmo2230* (E) and *lmo0596* (F) in wild type and Δ*sigB* taken 15 min after untreated (-) or treated (+) at pH 5.0. Gene expression was obtained from RT-qPCR and converted to Log_2_ gene expression relative to the wild type strain at time 0 min or untreated. Statistical analysis was performed using a paired student t test relative to the wild type untreated after 15 min, for RT-qPCR data, and untreated wild type at time 0 min, for acid challenge data (*, P value of <0.05; **, P value of <0.01; ***, P value of <0.001).

σ^B^ activation in response to sub-lethal acid stress was measured following exposure of exponential phase cultures to pH 5.0. To determine whether a sub-lethal acid exposure for 15 min influenced acid resistance, a survival experiment was performed at pH 2.5 to follow the resistance of bacteria that were either unadapted or pre-adapted at pH 5.0. There was a significant increase in acid resistance in the pre-adapted parental strain EGD-e (Fig 2D). As expected, the *sigB* deletion mutant was highly acid sensitive with no detectable survivors evident by 12.5 min post acid challenge (Fig 2D). At 15 min post adaptation there was a significant increase in the transcription of both *lmo2230* and *lmo0596* relative to the unadapted cultures (16-64-fold increase) in the wild type (Fig 2E and 2F). Together these data show that acid-induced σ^B^ activity can be measured at the transcriptional level and the role of σ^B^ in the acid resistant phenotype is evident under these conditions.

### RsbR1 paralogues are dispensable for acid signal transduction and acid adaptation

To investigate the role of the putative sensory component of the stressosome RsbR1 in transducing the acid signals into the σ^B^ activation pathway, we constructed a mutant strain where each of the three RsbR1 paralogues, RsbR2, RsbR3, and RsbR4 were deleted and the light sensing paralogue RsbL was inactivated by a mutation that produces a light-blind variant (C56A; (65)). The resulting genotype: Δ*rsbR2* Δ*rsbR3* Δ*rsbR4 rsbL*-C56A (42), is hereafter designated “*rsbR1*-only” for simplicity. When this mutant strain was subjected to the same acid exposure described above (pH 5.0), the transcription of the two σ^B^ reporter genes, *lmo2230* and *lmo0596*, was found to be strongly induced (Fig 3A and 3B) and there was a significant increase in acid resistance at pH 2.5, similar to that observed in the wild type (Fig 3C). The possibility that residual signal transduction activity was present in the mutated *rsbL-*C56A gene in the RsbR1-only strain was tested by measuring σ^B^ activation by sub-lethal acid exposure in a mutant background where all four paralogues were deleted (Δ*rsbR2* Δ*rsbR3* Δ*rsbR4* Δ*rsbL*) (Table 1). The behaviour of this strain was essentially identical to the RsbR1-only strain in terms of induced σ^B^ activity (S2A and S2B Fig) and acid resistance (S2C Fig). Thus, a strain lacking all known RsbR1 paralogues retains the ability to transduce acid signals via the stressosome and activate σ^B^ in response to acid adaptation at pH 5.0.

**Fig 3.**
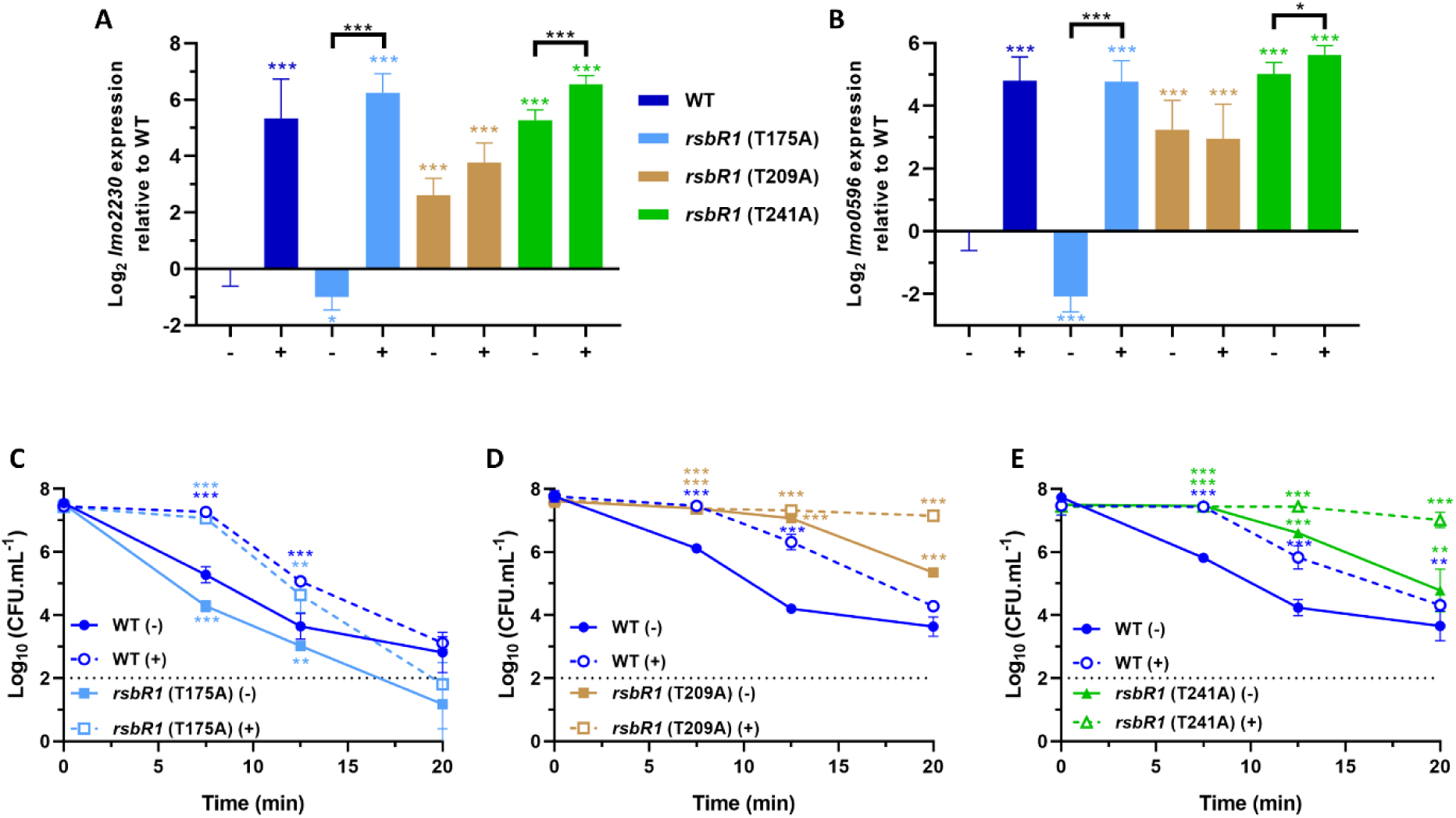
RsbR1 can act independently of its paralogues to transduce the low pH signals to activate σ^B^ and enhance acid tolerance. Expression of σ^B^-dependent genes (A) *lmo2230* and (B) *lmo0596*, obtained from mid-log phase cultures of wild type and *rsbR1*-only strains untreated (-) and treated (+) for 15 min in pH 5.0 at 37°C. Gene expression is expressed as Log_2_ relative gene expression. (C) Acid challenge of mid-log phase cultures treated in pH 5.0 for 15 min at 37°C, as previously described in Fig 2D. Survival data is expressed as Log_10_ (CFU.mL^-1^). The *rsbR1*-only genotype consists on the following mutations, Δ*rsbR2*; Δ*rsbR3*; Δ*rsbR4*; *rsbL* (C56A). Statistical analysis was performed using a paired student *t* test relative to the wild type untreated after 15 min, for RT-qPCR data, and untreated wild type at time 0 min, for acid challenge data (*, *P* value of <0.05; **, *P* value of <0.01; ***, *P* value of <0.001).

**Table 1.** Plasmids and strains used in this study.

| Plasmid or strain | Reference, Source |
| --- | --- |
| **Plasmids** |  |
| pMAD | (66) |
| pEX-K168::*rsbS* S56A | Eurofins Genomics |
| pMAD::*rsbR1* T175A | (42) |
| pMAD::*rsbR1* T209A | This study |
| pMAD::*rsbS* S56A | This study |
| pMAD::*rsbR1* T241A | This study |
| pMAD::*rsbT* N49A | (42) |
| pMAD::$\Delta rsbL$ | (67) |
| pIMK3 | (68) |
| pDNG7 (pIMK3-*P<sub>help::locOid::rsbR rsbS rsbT</sub>*) | This study |
| **Strains/Mutants** |  |
| *Escherichia coli* One Shot™ TOP10 | Invitrogen |
| *Listeria monocytogenes* EGD-e WT | K. Boor |
| *L. monocytogenes* EGD-e $\Delta sigB$ | (57) |
| *L. monocytogenes* EGD-e $\Delta prfA$ | (9) |
| *L. monocytogenes* EGD-e (*rsbL* C56A) | (65) |
| *L. monocytogenes* EGD-e (*rsbL* C56A Δ*rsbR2* Δ*rsbR3* Δ*rsbR4*) | (42) |
| *L. monocytogenes* EGD-e (Δ*rsbL* Δ*rsbR2* Δ*rsbR3* Δ*rsbR4*) | This study |
| *L. monocytogenes* EGD-e (*rsbL* C56A Δ*rsbR2* Δ*rsbR3* Δ*rsbR4* *rsbR1* T175A) | This study |
| *L. monocytogenes* EGD-e (*rsbL* C56A Δ*rsbR2* Δ*rsbR3* Δ*rsbR4* *rsbR1* T209A) | This study |
| *L. monocytogenes* EGD-e (*rsbL* C56A Δ*rsbR2* Δ*rsbR3* Δ*rsbR4* *rsbR1* T241A) | This study |
| *L. monocytogenes* EGD-e (*rsbL* C56A Δ*rsbR2* Δ*rsbR3* Δ*rsbR4*, *rsbS* S56A) | This study |
| *L. monocytogenes* EGD-e (*rsbR1* T175A) | (42) |
| *L. monocytogenes* EGD-e (*rsbR1* T209A) | This study |
| *L. monocytogenes* EGD-e (*rsbR1* T241A) | This study |
| *L. monocytogenes* EGD-e (*rsbS* S56A) | This study |
| *L. monocytogenes* EGD-e (*rsbT* N49A) | (42) |
| *L. monocytogenes* C12:C14 | (67) |
| **Complemented mutant strains** |  |
| *L. monocytogenes* EGD-e + pDNG7 | This study |
| *L. monocytogenes* EGD-e Δ*sigB* + pDNG7 | This study |
| *L. monocytogenes* EGD-e *rsbL* C56A Δ*rsbR2* Δ*rsbR3* Δ*rsbR4* + pDNG7 | This study |
| *L. monocytogenes* EGD-e *rsbR1* T175A + pDNG7 | This study |
| *L. monocytogenes* EGD-e *rsbR1* T209A + pDNG7 | This study |
| *L. monocytogenes* EGD-e *rsbR1* T241A + pDNG7 | This study |
| *L. monocytogenes* EGD-e *rsbS* S56A + pDNG7 | This study |
| *L. monocytogenes* EGD-e *rsbT* N49A + pDNG7 | This study |

### Role of stressosome phosphorylation in acid adaptation

A comparison of the STAS C-terminal domains of RsbR1 with the homologue RsbRA from *B. subtilis* reveals conserved threonine residues at positions 175, 209 and 241, respectively, of the *L. monocytogenes* protein, one of which (T175) is phosphorylated *in vivo* in both organisms ((42, 48); Fig 4A and 4B). To investigate the contribution of these phosphorylation sites in acid signal transduction these residues were substituted with alanine in the RsbR1-only background (Table 1) and the respective mutants subjected to the acid adaptation regime at pH 5.0. The *rsbR1*-T175A substitution did not prevent acid induction of σ^B^ activity (Fig 4A and B). In contrast the *rsbR1-*T209A substitution essentially abolished the acid induced transcription of the reporter genes and resulted in a significantly higher baseline transcription level compared to unadapted wild type bacteria (4 -16 fold higher; Fig 4A and 4B) (Christine Zielger, personal communication). Similarly, the *rsbR1*-T241A mutant also exhibited increased baseline transcription (Christine Zielger, personal communication), however the expression of these reporter genes was further increased by acid in this mutant. When the effects of these substitutions on acid resistance at pH 2.5 were measured, the *rsbR1*-T175A strain behaved similar to the wild type, showing an adaptive response to lethal acid challenge (Fig 4C). There was a small but significant decrease(∼ 1 log_10_ CFU.ml^-1^ between 7.5 and 20 min) in the acid resistance of the unadapted *rsbR1*-T175A mutant strain and this is correlated with a small reduction (∼ 1 fold; *p* = 0.016) in the transcription of the two reporter genes in the unadapted conditions compared to the wild type. The *rsbR1*-T209A and *rsbR1*-T241A mutants showed a marked increase (*p* < 0.001) in acid resistance compared to the wild type even in the unadapted culture (Fig 4D and 4E) and this correlated with the increased σ^B^ activity recorded by the reporter gene transcript levels (Fig 4A and 4B). We also examined *lmo2230* and *lmo0596* expression in genetic backgrounds with or without the RsbR1 paralogues when each of the three putative phosphorylation sites was separately mutated. Their expression pattern was not significantly influenced by the presence or absence of the RsbR1 paralogues (S3 and S4 Fig).

**Fig 4.**
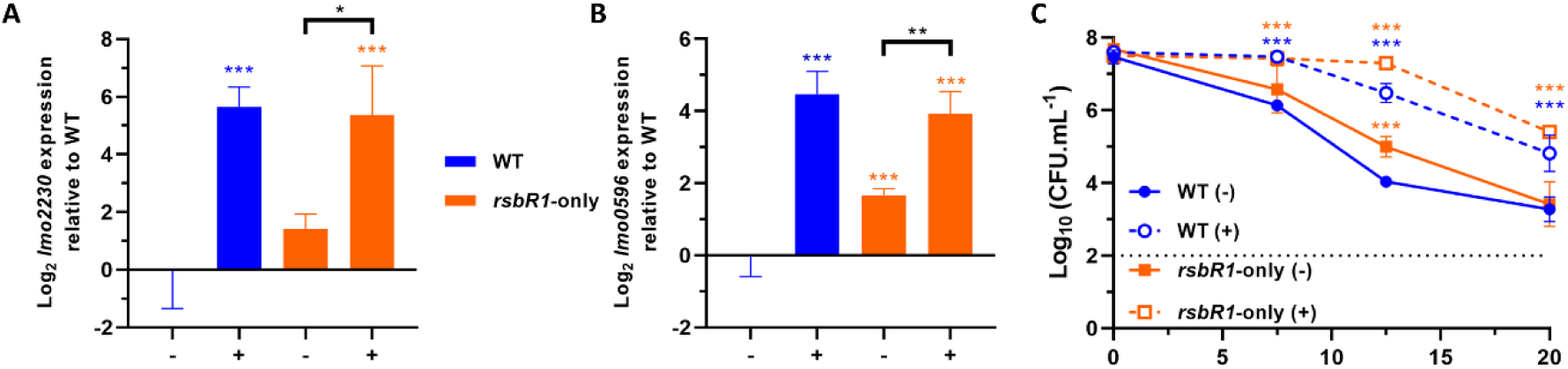
RsbR1 residues T209 and T241 are required for normal acid-mediated activation of σ^B^ and development of acid tolerance. Expression of σ^B^-dependent genes (A) *lmo2230* and (B) *lmo0596*, obtained from mid-log phase cultures of wild type and *rsbR1* (T175A), *rsbR1* (T209A) and *rsbR1* (T241A) strains untreated (-) and treated (+) for 15 min in pH 5.0 at 37°C. Gene expression is expressed as Log_2_ relative gene expression. Acid challenge of mid-log phase of (C) *rsbR1* (T175A), (D) *rsbR1* (T175A) and (E) *rsbR1* (T175A) cultures treated in pH 5.0 for 15 min at 37°C, as previously described in Fig 2D. Survival data is expressed as Log_10_ (CFU.mL^-1^). Statistical analysis was performed using a paired student *t* test relative to the wild type untreated after 15 min, for RT-qPCR data, and untreated wild type at time 0 min, for acid challenge data (*, *P* value of <0.05; **, *P* value of <0.01; ***, *P* value of <0.001).

The T209 residue of RsbR1 aligns with a conserved serine (S56) at the equivalent position in the STAS domain of *L. monocytogenes* RsbS (Fig 1B) and this RsbS residue is known to be important for signal transduction in the *B. subtilis* stressosome (49). We constructed a chromosomal allele of *rsbS* in the wild type background resulting in an RsbS S56A substitution (Table 1). The activation of σ^B^ during acid adaptation at pH 5.0 was completely abolished as indicated by the reporter genes. The transcription of these genes was also significantly lower (*p* = 0.0457 and <0.005 for *lmo2230* and *lmo0596*, respectively) than the wild type under unadapted conditions (Fig 5A and 5B). Adaptation at pH 5.0 to a lethal acid challenge (pH 2.5) was abolished in the *rsbS*-S56A mutant and the unadapted strain was also markedly more sensitive to acid that the wild type, showing no detectable survivors after 20 min (Fig 5C). Thus, the increased acid sensitivity of the *rsbS-*S56A mutant correlates with an inability to induce σ^B^ activity during the adaptation at pH 5.0, highlighting the importance of the S56 residue in the transduction of acid signals through the stressosome.

**Fig 5.**
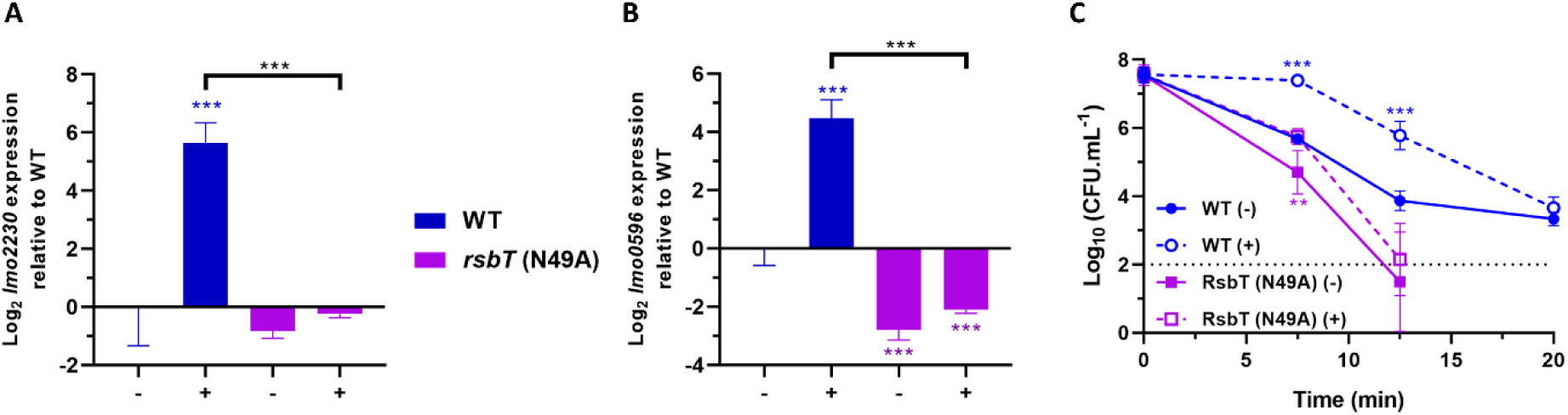
RsbS residue S56 is essential for the transduction of the low pH signal and activation of σ^B^. Expression of σ^B^-dependent genes (A) *lmo2230* and (B) *lmo0596*, obtained from mid-log phase cultures of wild type and *rsbS* (S56A) strains untreated (-) and treated (+) for 15 min in pH 5.0 at 37°C. Gene expression is represented as Log_2_ relative gene expression. (C) Acid challenge of mid-log phase cultures treated in pH 5.0 for 15 min at 37°C, as previously described in Fig 2D. Survival data is expressed as Log_10_ (CFU.mL^-1^). Statistical analysis was performed using a paired student *t* test relative to the wild type untreated after 15 min, for RT-qPCR data, and untreated wild type at time 0 min, for acid challenge data (*, *P* value of <0.05; **, *P* value of <0.01; ***, *P* value of <0.001).

To determine the role of stressosome phosphorylation in acid stress signal transduction, we used a strain where the kinase activity of RsbT was inactivated through mutation. The *rsbT*-N49A mutation was recently shown to abolish the kinase activity of RsbT (42). The acid induction of the two σ^B^ reporter genes at pH 5.0 was essentially eliminated in the *rsbT*-N49A background (Fig 6A and 6B) and this correlated with substantial decrease in acid resistance at pH 2.5 (Fig 6C).

**Fig 6.**
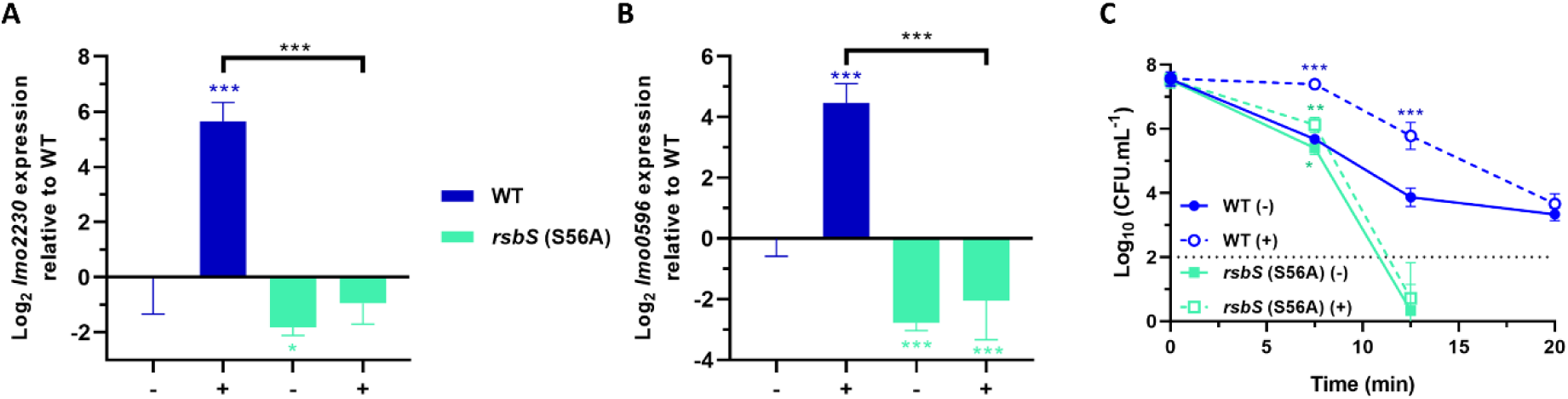
The kinase activity of RsbT is essential for acid sensing by the stressosome. Expression of σ^B^-dependent genes (A) *lmo2230* and (B) *lmo0596*, obtained from mid-log phase cultures of wild type and *rsbT* (N49A) strains untreated (-) and treated (+) for 15 min in pH 5.0 at 37°C. Gene expression is represented as Log_2_ relative gene expression. (C) Acid challenge of mid-log phase cultures treated in pH 5.0 for 15 min at 37°C, as previously described in Fig 2D. Survival data is expressed as Log_10_ (CFU.mL^-1^). Statistical analysis was performed using a paired student *t* test relative to the wild type untreated after 15 min, for RT-qPCR data, and untreated wild type at time 0 min, for acid challenge data (*, *P* value of <0.05; **, *P* value of <0.01; ***, *P* value of <0.001).

To confirm that the mutations introduced into *rsbR1*, *rsbS* and *rsbT* were responsible for the observed effects on acid survival, we provided a functional copy *in trans*. The single mutant strains were transformed with an IPTG inducible vector (pDNG7) bearing the parental *rsbR1-rsbS-rsbT* open reading frames (Table 1), were grown to stationary phase, where σ^B^ is highly active, and challenged in BHI at pH 2.5 at 37°C. The wild type strain showed further increase in acid tolerance when IPTG was added to the medium (S6A and S6B Fig). Interestingly, the acid tolerance of the *rsbR1* T209A and T241A strains, that had elevated baseline σ^B^ activity, showed no further increase, suggesting that the complementation with native stressosome is unable to outcompete the mutated RsbR1 and attenuate σ^B^ activity. Finally, the complementation of the stressosome inactive strains, *rsbS* S56A and *rsbT* N49A, successfully restored their phenotypes but not for the Δ*sigB* strain (S6A-S6C Fig), showing that the provision of native stressosome *in trans* can restore the signal transduction in these strains.

To investigate the effects of the *rsbR1* (T175A, T209A and T241A), *rsbS* (S56A) and rsbT (N49A) mutations on stressosome phosphorylation following sub-lethal acid treatment, we determined the phosphorylation patterns of RsbR1 and RsbS using the Phos-Tag electrophoresis system. A single RsbR1 band, probably corresponding to the unphosphorylated isoform, was observed in the *rsbT*-N49A kinase mutant. In contrast, two bands with reduced electrophoretic mobility were apparent in the wild type, presumed to correspond to different phosphorylated isoforms of higher molecular weight (Fig 7A and S5 Fig). The *rsbR1-*T175A and *rsbR1-*T209A mutations each resulted in the loss of one of the upper bands, suggesting that each band might be represent a different singly phosphorylated isoform. It was not clear whether this method could adequately resolve a doubly phosphorylated form of RsbR1. The acid adaptation at pH 5.0 had minimal effects on the phosphorylation patterns detected, although there was some evidence in the wild type, *rsbR1-*T175A and *rsbS-*S56A backgrounds that T209 was slightly more phosphorylated following acid exposure (Fig 7A and 7B). Anti-RsbS blotting confirmed the disappearance of the phosphorylated isoform (upper band) of RsbS in the *rsbS-*S56A and *rsbT-*N49A backgrounds (Fig 7B). The specificity of the anti-RsbS antibodies was confirmed using a Δ*rsbR1* mutant that is polar and abolishes RsbS expression (42). Overall these data confirmed the loss of phosphorylation of RsbR1 and RsbS in the site directed mutant strains, thereby supporting the conclusion that phosphorylation of RsbR1 (at residue T209) and RsbS (at residue S56) is necessary for transducing acid stress signals through the stressosome.

**Fig 7.**
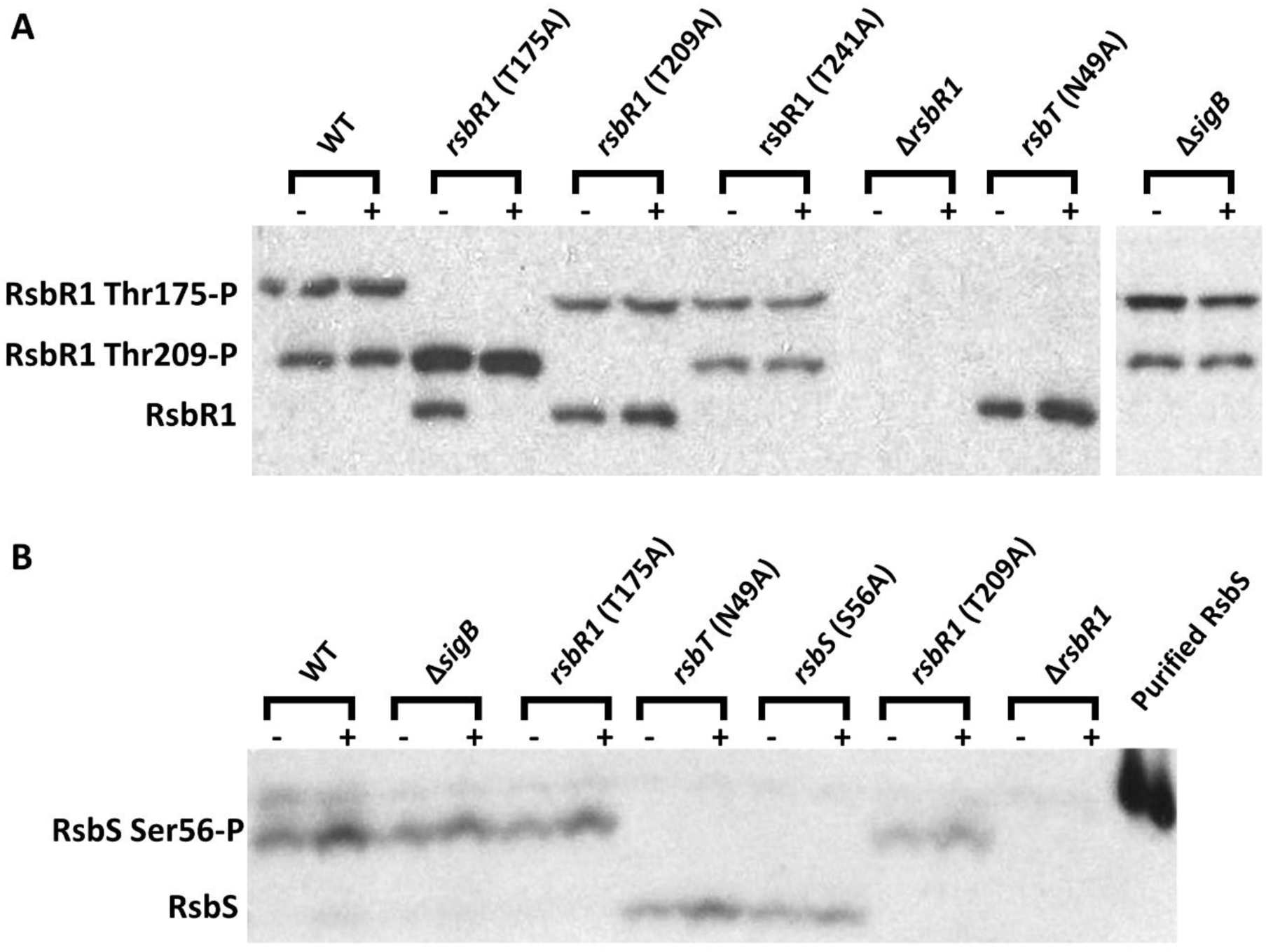
Phosphorylation of RbsR1 on residues T175, T209 and RsbS on residue S56 requires the kinase RsbT. Western blots probed with (A) anti-RsbR1 and (B) anti-RsbS antibodies. Wild type and all mutant strains were grown to mid-log phase at 37°C in BHI and then untreated (-) or treated (+) at pH 5.0 for 15 min. Total protein extracted was separated in Phos-tag gels. The lower bands correspond to non-phosphorylated isoforms of RsbR1 or RsbS in the respective blots. The middle bands in anti-RsbR1 blots correspond to mono-phosphorylated RsbR1 at the T209 and the higher band corresponds to mono-phosphorylated RsbR1 at T175. The higher band on the RsbS blot corresponds to RsbS phosphorylated on S56. The Δ*rsbR1* mutant extracts were included as negative controls since RsbR1 and RsbS are not expressed in this strain.

Finally, the phosphorylation patterns of RsbR1 were not affected in the *rsbR1*-T241A mutant. This mutant has previously been shown to have increased baseline transcription of *lmo2230* and *inlA* similar to *rsbR1*-T209A (Christine Zielger, personal communication), which together with the structural data led the authors to suggest a putative interaction between T209-P and T241 in RsbR1. The finding that substitution of either of these residues produces a similar effect on acid signalling (Fig 4A and 4B) and acid resistance (Fig 4D and 4E) provides genetic support for this putative regulatory interaction.

### Acid-induced virulence gene expression requires functional stressosome signal transduction

There are several lines of evidence that σ^B^ plays an important role in transcription during the gastrointestinal stage of the infectious cycle (9,69,70). We investigated whether the invasion genes *inlA* and *inlB*, required for penetration across the intestinal barrier, and known to be partly under σ^B^ control (13, 14), might be induced in response to the low pH signals found in the GI tract. Using the same acid adaptation regime described above (pH 5.0 for 15 min) we found that both *inlA* and *inlB* were strongly induced (∼ 16-fold) in the wild type but not in a *sigB* mutant (Fig 8A and8 B). Since both genes belong to one operon, the shared pattern of expression was not surprising. Next the role of the stressosome in this acid induced regulation was investigated. A strain carrying RsbR1 but none of its four paralogues (RsbR1-only) showed a very similar pattern of *inlAB* transcription to the wild type. Mutations abolishing the kinase activity of RsbT (*rsbT*-N49A) or the phosphorylation of RsbS (*rsbS-*S56A) eliminated the acid-induced transcription of the *inlAB* operon (Fig 8A and B). Mutations in *rsbR1* produced a less pronounced effect on *inlAB* induction by acid, with the *rsbR1-*T175A mutant showing a very similar response to the wild type and the *rsbR1-*T209A mutant showing a significantly higher level of *inlAB* transcription in the untreated condition (Fig 8A and 8B). Together these data strongly suggest that RsbR1 is sufficient to sense and transduce acid stress signals into the stressosome and show that this is necessary for acid induced regulation of the virulence invasion genes *inlA* and *inlB*.

**Fig 8.**
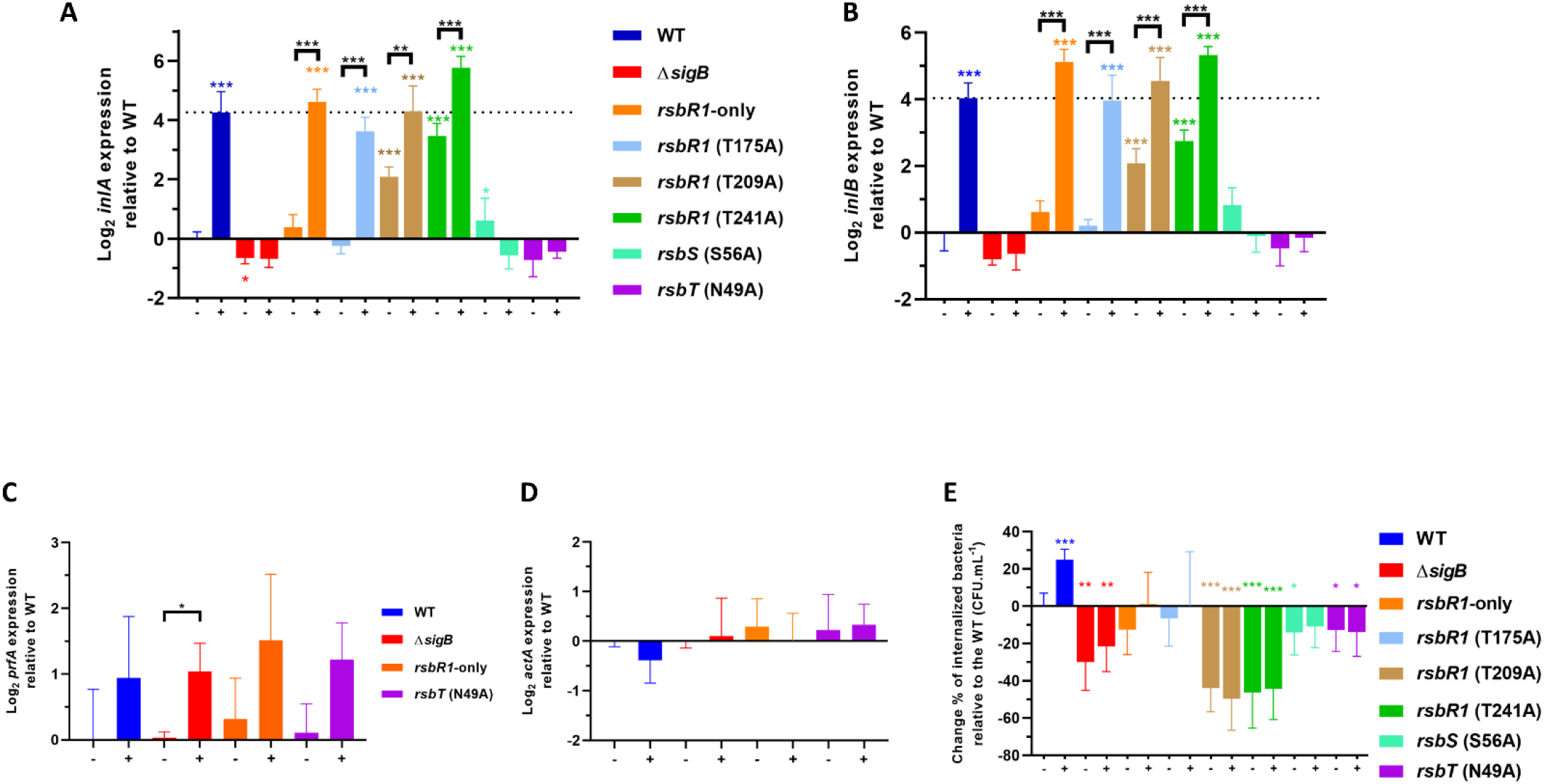
Acid sensing via the stressosome triggers upregulation of the *inlAB* invasion operon and influences Caco-2 cell invasion without affecting PrfA activity. (A) *inlA,* (B) *inlB*, (C) *prfA* and (D) *actA* expression was obtained from mid-log phase cultures of strains used in this study. Cultures were untreated (-) and treated (+) in pH 5.0 for 15 min at 37°C. (E) Caco-2 invasion experiments performed on untreated (-) and treated (+) cultures in pH 5.0 for 15 min. *L. monocytogenes* wild type and mutant strains were in contact with the Caco-2 epithelial cells for 1 h and extracellular bacteria were eliminated with extensive washing and the supplementation of 50 µg.mL^-1^ of gentamicin to fresh tissue culture medium. Quantification of internalized *L. monocytogenes* was obtained from lysed Caco-2 cells 1 h after the gentamicin treatment. The untreated wild type strain was arbitrary converted to 0% and all samples were normalized to percentage of CFU.mL^-1^ relative to the untreated wild type. Statistical analysis was performed using a paired student *t* test relative to the wild type untreated after 15 min (*, P value of <0.05; **, P value of <0.01; ***, P value of <0.001).

The possibility that induction of *inlAB* by acid could involve the virulence gene regulator PrfA was examined by looking at *prfA* transcription and the PrfA-dependent transcription of *actA* under the same experimental conditions. The *prfA* transcript increased slightly (∼2-fold) following acid treatment, as has been reported by others (17). This increase was evident in a mutant lacking σ^B^ and all the strains carrying RsbR1, RsbS and RsbT mutations (Fig 8C) and was therefore independent of σ^B^ and stressosome function. This small increase in *prfA* transcript levels did not affect PrfA activity since the levels of the highly PrfA-dependent (but not σ^B^-dependent) transcript *actA* were not affected by acid treatment (Fig 8C-8D). Together these data confirm that the involvement of σ^B^ in the acid-induced transcription of *inlAB* was direct and did not involve an indirect effect on PrfA.

Finally, we investigated whether the acid-induced activation of σ^B^ via the stressosome affected host cell invasion. Compared to the untreated wild type bacteria, the acid-exposed wild type bacteria displayed an increase in Caco-2 cell uptake of approximately 25% in the invasion of Caco-2 epithelial cells (Fig 8E). Mutants lacking *sigB* or carrying any of the stressosome alleles all showed a reduced invasion under the untreated conditions and the presence of acid failed to stimulate invasion. Surprisingly, even those strains that showed increased *inlAB* transcription (RsbR1 T209A, RsbR1 T241A) showed a reduced invasion rate compared to the wild type parental strain (Fig 8E). It is possible that post-transcriptional control of *inlAB* expression influenced the invasion rate, but further investigation of this phenomenon was beyond the scope of this study. Overall, the data showed that mutations that blocked acid induced σ^B^ activation also affected the acid stimulated increase in host cell invasion.

## Discussion

In this study, we sought to understand the signal transduction mechanisms that allow the stressosome of *L. monocytogenes* to transduce acid stress signals leading to activation of σ^B^ and the GSR. While it has been known for over two decades that σ^B^ plays a critical role in acid stress tolerance in this pathogen (18, 71) nothing was known about how, or indeed whether, acid stress influenced σ^B^ activity. Here we show that the *L. monocytogenes* stressosome is responsible for the rapid transduction of the environmental low pH signal. Specifically RsbR1 in conjunction with RsbS and RsbT, independently of the other four RsbR1 paralogues, are sufficient to initiate the downstream signal transduction events leading to σ^B^ activation. The kinase activity of RsbT and phosphorylation of RsbS on residue S56 are essential for transduction of the acid stress signal. The activation of σ^B^ by mild acidic stress (pH 5) results in the upregulation of σ^B^-dependent genes and produces an increase in acid resistance to lethal acid pH challenge.

Although the overall composition of the *L. monocytogenes* stressosome, which comprises RsbR1 and its paralogues (RsbL, RsbR2, RsbR3), RsbS and RsbT, is now fairly well understood (37,42,44), the stoichiometry of individual stressosomes and heterogeneity of stressosome composition within cells and populations is still unknown. The outward facing turrets on the stressosome, derived from the N-terminal domains of RsbR1/RsbRA and paralogues, have been suggested to be the most likely sensory domains, although the nature of the signals detected has in most cases been elusive. As postulated by Murray and colleagues, the non-haem globin domains, found in most RsbR1/RsbRA-like proteins, may transiently interact with small signal molecules (72), but evidence for these has not been forthcoming. There may be some redundancy in the sensory capacity of RsbR1/RsbRA-like proteins since studies on the *B. subtilis* stressosome have shown that different RsbR paralogues can respond to ethanol stress (49, 73). Here we sought to establish whether RsbR1 was capable of sensing and transducing acid stress signals via the stressosome. While it was not possible to delete *rsbR1* without affecting the expression of the downstream genes (*rsbS, rsbT, rsbU;* (42)) it was possible to construct a strain lacking functional versions of all four paralogues (RsbL, RsbR2, RsbR3, RsbR4) and this mutant displayed a near-wild type activation of σ^B^ in response to acid stress (Fig 3A and 3B). The results show that RsbR1 is capable of sensing and transducing acid stress, independently of its paralogues, although the precise nature of the signals detected will require further research.

From studies on the non-pathogenic bacterium *B. subtilis* we have learned that the stressosome is a dynamic protein complex whose activation depends on phosphorylation of residues in RsbRA and RsbS (48, 51), although acid stress has not been investigated as a stressor in this organism. In *L. monocytogenes,* RsbR1 is phosphorylated on residues T175 (74) and T209, but the phosphorylation sites are not conserved in the RsbR1 paralogues (Fig 1B). In this study, we confirm the phosphorylation of RsbR1 T209 and show that loss of this phosphorylation leads to derepressed σ^B^ activity under no-stress conditions and loss of responsiveness to acid. The finding that a substitution of RsbR1 residue T241 with alanine produces a similar phenotype provides further evidence that these two residues may interact, as suggested by the structure predicted recently by cryoEM (Christine Zielger, personal communication).

Although a detailed mechanistic model accounting for signal transduction through the stressosome is not possible at this time, we propose the following working model to aid further experimental work on the system. The outward-facing dimeric N-terminal turret domains of RsbR1 most likely function in signal detection (75), although the nature of that signal remains to be determined. The acid stress signal could either be a change in the local proton concentration or some secondary consequence of acidification. The signal is likely to produce a conformation change of the N-terminal domains that propagates into the alpha-helical linker domains and into the STAS domains, resulting in the release of RsbT and consequently activation of σ^B^ via the downstream partner switching module (39, 76). We propose that the release of RsbT from the stressosome likely requires the phosphorylation of RsbS on S56, since acid sensing is abolished in the RsbS S56A variant and in the kinase-inactive mutant (Figs 5 and 7B). In this regard it is noteworthy that substitutions in *B. subtilis* RsbS (S59D) and *Moorella thermoacetica* MtS (S58D; MtS is a homologue of RsbS) abolish the interaction with the cognate kinase (77, 78). Furthermore, these mutations lock the stressosome in an active state, resulting in constitutive σ^B^ activity, which is deleterious for the bacteria (51,78,79).

The role of RsbR1 phosphorylation on residues T175 and T209 in modulating the response to acid is less clear but we speculate that they may influence the propagation of the signal that results in phosphorylation of RsbS. Altering T209 to alanine increased the background σ^B^ activity in the absence of acid stress and essentially abolished the responsiveness of the system to acid, while the T175A substitution did not affect acid sensing (Fig 4). It is possible that they serve to modulate the rate of RsbS phosphorylation by RsbT thereby affecting the overall kinetics of the response, but a more time resolved analysis will be needed to investigate this. The finding that the phosphorylation state of RsbR1 and RsbS doesn’t change much in response to osmotic (42) or acid stress (Fig 7A and 7B) is apparently in contradiction with the finding that phosphorylation is necessary for signal transduction, as evidenced by the absence of signal transduction in a kinase deficient mutant (Fig 6; (42)). One possibility we have not ruled out is that the method we used to determine the phosphorylation state of RsbR1 and RsbS is not sufficiently robust to exclude the possibility of phosphorylation during the protein extraction procedure (which includes a boiling step). Phosphomimetic substitutions that add negatively charged residues to the phosphorylatable sites in RsbR1 and RsbS will help to further clarify this issue.

While the phosphorylation undoubtedly plays a pivotal role in the activation of the stressosome, the precise mechanisms that result in release of RsbT are unknown at present. It has been suggested that conformational changes may occur at the STAS domains of RsbR1 and RsbS, prompting the release of RsbT from the stressosome core (39). A recent study investigating the *L. monocytogenes* stressosome structure found T241 residing in RsbR1 α3 helix, described as “flexible loop”, which is located in close proximity to the T209 phosphorylation site (44). Moreover, when phosphorylated, T209 forms a hydrogen bound with T241 (Christine Zielger, personal communication). Given that amino acid substitutions to alanine at both of T209 and T241 lead to constitutive σ^B^ activity (Fig 4A and 4B and S3C-S3F Fig), we speculate that the phosphorylation of T209 and its subsequent interaction with T241 likely determines the relative position of this flexible loop within the stressosome. We speculate that the loop’s position may influence the magnitude of the signal transduced by the stressosome, perhaps by affecting the propagation of the signal from one protomer to another. Further structural and genetic studies will be needed to fully elucidate the role phosphorylation in signal transduction. The role of the putative phosphatase RsbX in resetting the sensing–ready state of the stressosome has recently been suggested (54) but further work will be needed to elucidate the details of precisely how it interacts with the stressosome.

Multiple studies have shown that pre-treating *L. monocytogenes* with mild acidic pH increases its invasion towards Caco-2 cells (15–17) and this correlates well with increased *inlA* expression under these conditions (15,17,23). Our study shows that the σ^B^-dependent induction of the *inlAB* operon under mild acidic conditions is dependent on the stressosome’s sensory capacity (Fig 8A and 8B). In addition, we showed that the enhancement of *L. monocytogenes* EGD-e invasion is critically dependent on the normal function of stressosome sensing properties and a proper σ^B^ upregulation, as either lower or higher baseline activity abrogates *L. monocytogenes* capacity to invade Caco-2 cells (Fig 8E). Interestingly, the high baseline σ^B^ activity produced in the RsbR1 T209A and T241A strains correlated with a decrease in invasion, even though transcription of the *inlAB* operon was increased in this genetic background. This result demonstrates clearly that host cell invasion does not solely depend on the transcription of invasion genes, and further studies will be required to determine if this effect is due to some unknown post-transcriptional effect on *inlAB* expression. Indeed post-transcriptional control of *inlA* expression has been reported before (80). Overall, our results support the idea that σ^B^ activity during the early stages of the infection cycle plays crucial role in priming the pathogen for host cell invasion and that the stressosome participates by helping to integrate pH-related signals from the environment of GI.

Overall, this study has established that the stressosome plays an active role in sensing and responding to acid stress by modulating the expression of the GSR. The signal transduction mechanism requires the kinase activity of RsbT and the phosphorylation of S56 on RsbS. In RsbR1, both T209 and T241 constrain σ^B^ activation in no-stress conditions and are necessary for a normal response to acid stress, while T175 appears to be dispensable for acid sensing. This signal transduction pathway has a demonstrable role in the development of acid resistance, likely influencing the survival of this pathogen in the food chain and in the acidic conditions encountered within the host gastrointestinal tract. This pathway likely influences the progression of the infectious cycle during listeriosis through the impact on the expression of virulence factors that influence host cell invasion in the GI tract.

## Material and Methods

### Bacterial strains, plasmids, and primers

*L. monocytogenes* EGD-e (serovar 1/2a) and *E. coli* TOP10 strains, plasmids and primers used in this study are listed in Table 1 and Table 2. Strains were grown in BHI broth or agar (LabM, UK) at 37°C with constant shaking at 150 rpm.min^-1^ at initial neutral pH ∼ 7.4, unless otherwise specified. Cells were grown for 16 h until stationary phase was reached or further diluted in fresh BHI to an initial OD_600_ = 0.05 and further allowed to grow until mid-log phase (OD_600_ = 0.4). The following antibiotics were added to the media to a final concentration when required: erythromycin (Ery) at 5 µg.ml^-1^ for *L. monocytogenes* strains, ampicillin (Amp) at 100 µg.ml^-1^ for *E. coli* strains and kanamycin (Km) 50 µg.ml^-1^ for both microorganisms.

**Table 2.**
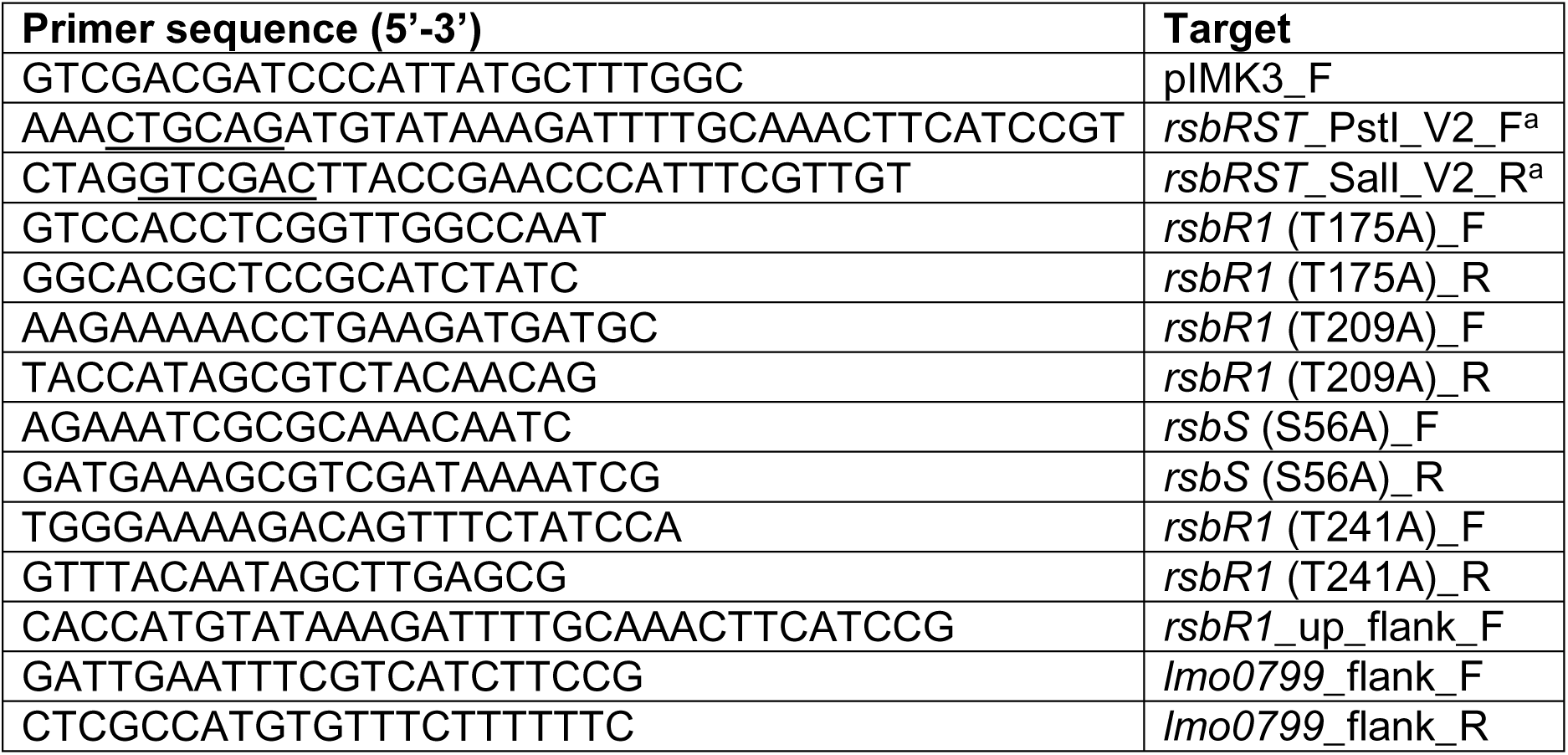
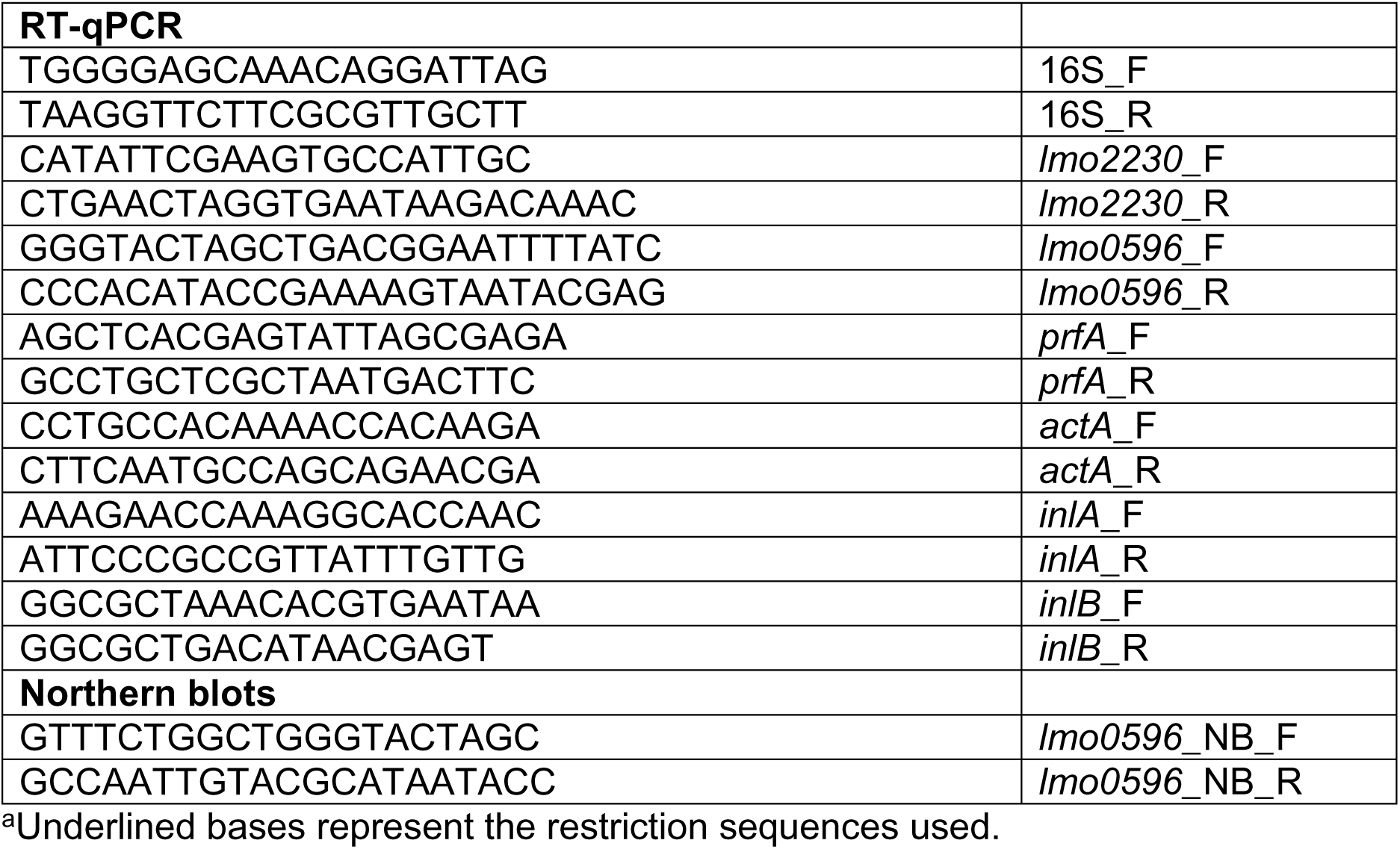
Primers used in this study.

### Construction of genetically modified *L. monocytogenes*

The gene *rsbS* Serine-to-Alanine in the codon 56. The codon 56 TCA (Serine) in *rsbS* was changed to GCT (Alanine). In both cases silent mutations were added the two adjunct codons in order to discern mutant from WT codons by PCR during mutagenesis while taking in consideration the codon frequencies in *L. monocytogenes* EGD-e strain (*rsbS* WT -ATTGAT<u>TCA</u>TTTATT- to *rsbS* S56A - ATCGAC<u>GCT</u>TTCATC-). The mutagenic sequences, each with a total length of 612 bp, including SalI and BamHI restriction sites in each edges, were artificially synthetized in the vector pEX-K168::*rsbS* (S56A) (Eurofins Genomics, Germany). The mutagenic sequences were subsequently cloned into shuttle vector pMAD, originating pMAD::*rsbS* (S56A). *L. monocytogenes* electrocompetent cells were created as previously described (68). The electrocompetent *L. monocytogenes* WT strain was transformed with pMAD::*rsbS* (S56A). The quadruple mutant strain (*lmo0799* (C56A); Δ*lmo0161*; Δ*lmo1642*; Δ*lmo1842*) was further separately transformed with pMAD::Δ*lmo0799,* pMAD::*rsbR1* (T175A), pMAD::*rsbR1* (T209A) and pMAD::*rsbS* (S56A). Electroporated cells were plated in BHI supplemented with Ery. Chromosomal integration and subsequent excision was achieved through a two-step recombination as previously described (65). cPCR with the primers *rsbR1*_upflank_F paired with either *rsbR1* (T175A)_R, *rsbR1* (T209A)_R, *rsbR1* (T241A)_R and *rsbS* (S56A)_R, were used to identify the chromosomal mutation in the respective genes *rsbR1*, *rsbS* and *rsbT*, while the deletion of *lmo0799* was verified with primers *lmo0799*_flank_F and *lmo0799*_flank_R. All genomic DNA from each mutant strains were whole genome sequence to confirm the chromosomal mutation and identify additional secondary mutations.

### Growth kinetics

Stationary-phase cells of the *L. monocytogenes* EGD-e WT and Δ*sigB* were diluted and adjusted to an initial OD_600_ of 0.05 and grown in fresh BHI at 37°C. OD_600_ measurements were taken every hour for 10 h. Three biological replicates were made.

### Acid shock treatment in *L. monocytogenes*

Stationary phase cultures grown at 37°C were diluted to an initial OD_600_ of 0.05 in fresh BHI and allowed to grow at 37°C until mid-log phase was reached (OD_600_ = 0.4). Cultures were either acidified or not until pH 5.0 with HCl 5 M and further incubated for 15 min at 37°C. Cultures were diluted in 1:10 of BHI previously acidified to pH 2.5 with HCl 5 M. Samples were taken at 0, 7.5, 12.5 and 20 min, serial diluted from 10^-2^ to 10^-7^ in PBS and plated in BHI agar. Plates were then incubated at 37°C for 24 hours and colonies were counted. At least three biological replicates were made.

### RNA extraction and RT-qPCR

Cultures of *L. monocytogenes* EGD-e and its isogenic mutants were grown until mid-log phase and subjected to acid treatment for 15 min as previous described. Cultures were diluted in RNAlater^TM^ (Sigma, Germany) at a 1:5 ratio to stop the transcription. Total RNA was extracted using an RNeasy® minikit (Qiagen, Nertherlands) according to the manufacturer’s recommendations. Cells were disrupted by bead beating twice in FastPrep-24 at a speed of 6 m.s^-1^ for 40 s. DNA was digested with Turbo DNA-free (Invitrogen, UK) according to the manufacturer’s recommendations. The RNA integrity was verified by electrophoresis in 0.7% (w/v) agarose gels. Synthetize of cDNA was performed with SuperScript™ III First-Strand Synthesis System (Invitrogen) according to the manufacturer’s recommendations. cDNA was quantified using NanoDrop 2000c (ThermoFisher Scientific, UK) and diluted to a final concentration of 7 ng.ml^-1^. RT-qPCR was performed using a QuantiTect SYBR® Green PCR kit (Qiagen) and primers for the target genes (Table 2). Primer efficiency for 16S, *lmo2230*, *lmo0596*, *inlA*, *inlB*, *prfA* and *actA* were previously tested using cDNA. Samples were analysed on LightCycler® 480 system (Roche) with the following parameters: 95°C for 15 min; 45 cycles of 15 s at 95°C, 15 s at 53°C, and 30 s at 72°C; a melting curve drawn for 5 s at 95°C and 1 min at 55°C, followed by increases of 0.11°C.s^-1^ until 95°C was reached; and cooling for 30 s at 40°C. Cycle quantification values were calculated by using LightCycler® 480 software version 1.5.1 (Roche, Switzerland) and the Pfaffl relative expression formula (81, 82). The expression of 16S rRNA was used as a reference gene. Results are expressed as Log_2_ relative expression ratios normalized against the expression of *L. monocytogenes* WT strain in the absence of stress. At least three independent biological replicates were performed.

### Time-lapse of σ^B^ response towards mild acidic pH

Stationary cultures of *L. monocytogenes* WT and Δ*sigB* were grown at 37°C were diluted to an initial OD_600_ of 0.05 in fresh BHI and allowed to grow at 37°C until mid-log phase was reached (OD_600_ = 0.4). Cultures were then acidified until pH 5.0 with HCl 5 M and further incubated up to 30 min. Samples were taken at 0, 5, 15 and 30 min post acidification and RT-qPCR were performed. At least three biological replicates were made.

### Caco-2 cell invasion assay

Caco-2 cell line human colon adenocarcinoma (Merck, UK) were cultivated in DMEM (Sigma) supplemented with 10% (v/v) FBS (Sigma), 1x MEM non-essential Amino acids (Gibco, UK) and Pen-Strep (Merck) at 37°C, in 5% CO_2_ and at 90-95% humidity. 5 x 10^4^ cells of non-differentiated Caco-2 (passages 12 to 15) were seeded 48 h prior infection in a 24 well-plate. *L. monocytogenes* strains were separately grown until mid-log phase (OD_600_ = 0.4) and acid shocked at pH 5.0 for 15 min at 37°C and washed twice in DPBS and adjusted to a multiplicity of infection of 100 in DMEM without antibiotics. Caco-2 cells were covered with DMEM containing *L. monocytogenes* strains. The bacteria were in contact with the eukaryotic cells for 1 h and subsequently washed twice with PBS and covered with 500 µL of DMEM supplemented with 50 µg.mL^-1^ of gentamicin for 1 h. Subsequently, the wells were washed twice with DPBS and the Caco-2 cells were lysed with 1 mL of cold PBS supplemented with 0.1% (v/v) Triton X-100. Intracellular bacteria was determinate by plating serial diluted lysates in BHI and incubated for 24 h at 37°C. Results were calculated with the difference between the CFU.mL^-1^ counts of the untreated wild type strain and each respective mutant.

### Identification of phosphorylated isoforms of RsbR1 and RsbS

Cultures and protein extraction were performed as previously described by others (42) with some minor changes. *L. monocytogenes* WT and its isogenic mutants were acid shocked cultures before being boiled for 20 s at 100°C to stop the metabolism before harvesting the cells by centrifugation. Total protein standardization was confirmed by SDS-PAGE (S5 Fig).

### Northern-blots probed for *lmo0596*

Total RNA was isolated from *L. monocytogenes* WT, Δ*sigB*, Δ*prfA* and the transposon mutant C14:C12, containing a frameshift in *rsbS* open reading frame, as previously described by others (83) with some modifications. WT and mutants strains were allowed to grow until late-log phase (OD_600_ = 0.8) and acid shocked before transcription was stopped. The DNA probe for *lmo0596* was generated as previously described (67).

### Validation of *lmo0596* as a σ^B^-dependent gene under mild low pH stress

Late-log phase cultures (OD_600_ = 0.8) of *L. monocytogenes* WT, Δ*sigB* and Δ*lmo0596* were acid shocked at pH 5.0, while WT and Δ*sigB* strains were treated with a range of pH’s (non-acidified pH 7.0, 6.0, 5.5, 5.0 and 4.5). Total RNA extraction, cDNA synthesis and RT-qPCR were performed as described above.

### Genetic complementation of *L. monocytogenes* mutant strains, acid tolerance and Congo Red phenotyping

Genetic complementation was achieved by amplifying *rsbR1*-*rsbS*-*rsbT* open reading frames using the primers *rsbRST*_PstI_V2_F and *rsbRST*_SalI_V2_R. The obtain amplicon and the IPTG inducible vector, pIMK3, were separately digested using PstI and SalI and ligated afterwards. The resulting plasmid, pDNG7, was transformed into *E. coli* TOP10 competent cells. The pDNG7 vector was transformed into electrocompetent *L. monocytogenes* strains and plated into BHI supplemented with Kanamycin 50 µg.ml^-1^. Complementation was verified using the primers pIMK3_F and r*sbRST*_SalI_V2_R on the obtained colonies. Obtained complemented strains were grown at 37°C for 16 h in BHI supplemented or not with 1 mM of IPTG (Sigma) until stationary phase was reached. Strains were challenged in BHI (pH 2.5) acidified with HCl 5 M and samples were taken at 0 and 30 min post challenge, serial diluted in PBS and plated in BHI agar plates. 10 µL of stationary phase cultures of the complemented strains were spotted in BHI agar plates supplemented with Congo Red (Sigma) 25 µg.mL^-1^ and IPTG 1 mM. Plates were incubated at 37°C then transferred to 30°C for 3 days. At least two biological replicates were made.

### Whole-genome sequencing and Single Nucleotide Polymorphism (SNP) analysis

The gDNA of all mutants strains constructed in this study was extracted using DNeasy® Blood and Tissue kit (Qiagen) according to the manufacturer recommendations. The obtained genomic material was analysed via Illumina sequencing by MicrobesNG (Birmingham, UK) and the resulting trimmed reads were analysed using Breseq software (84) to identify additional SNP in the mutant strains chromosome. The nucleotide sequence of *L. monocytogenes* EGD-e chromosome (NCBI RefSeq accession no. NC_003210.1) was used as reference in this analysis.

### Statistical analysis

All statistical analysis, student’s *t* test were performed using GraphPad Prism 8.

## Acknowledgments

We are grateful to Christine Zielger (University of Regensburg) and her colleagues for helpful discussions and for sharing results with us prior to their publication. We thank members of the PATHSENSE consortium for helpful discussions and suggestions during the course of this research.

## Supporting information

**S1 Fig.**
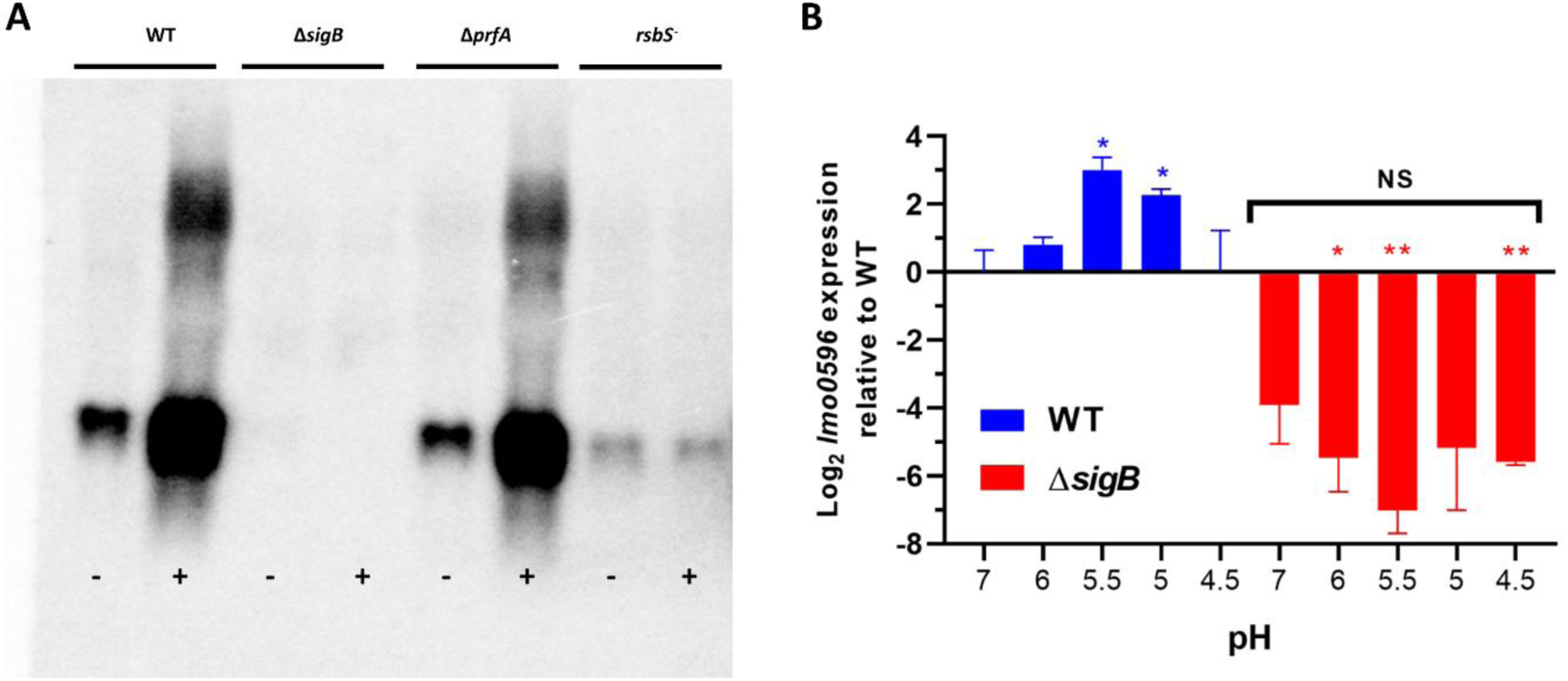
The transcription of *lmo0596* is induced by low pH in a highly σ^B^-dependent manner. Cultures of wild type, Δ*sigB*, Δ*prfA* and *rsbS*^-^ (formerly C12:C14) strains were grown to late-log phase (OD_600_ = 0.8) and untreated (-) and treated (+) samples in pH 5.0 or a range of pH (7.0 to 4.5) for 15 min. Total RNA was extracted as described in material and methods. *rsbS*^-^ mutant consists on a transposon strain which also carries a frameshift in *rsbS* resulting in the premature stop codon and a polar effect on RsbT translation (57). (A) Northern blots probed for *lmo0596*. (B) RT-qPCR results for *lmo0596* expression in wild type and Δ*sigB* strains in BHI adjusted to a pH range (7, 6, 5.5, 5 and 4.5). Gene expression obtained by RT-qPCR is expressed as Log_2_ relative gene expression. Statistical analysis was performed using a paired student *t* test relative to the untreated wild type after 15 min (NS = not significant; *, P value of <0.05; **, P value of <0.01; ***, P value of <0.001).

**S2 Fig.**
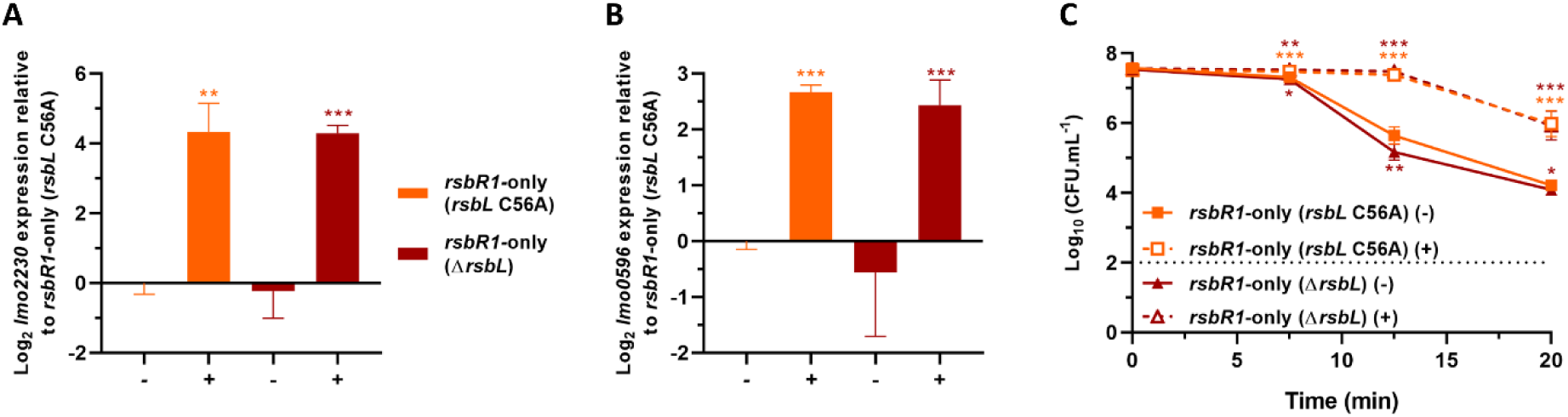
RsbL does not play a role in the stressosome acid sensing mechanism. Expression of σ^B^-dependent genes (A) *lmo2230* and (B) *lmo0596*, obtained from mid-log phase cultures of *rsbR1*-only (*rsbL* C56A) and *rsbR1*-only (Δ*rsbL*) strains untreated (-) and treated (+) for 15 min in pH 5.0 at 37°C. (C) Acid challenge of mid-log phase cultures treated in pH 5.0 for 15 min at 37°C, as previously described in Fig. 2D. Survival data is expressed as Log_10_ (CFU.mL^-1^). The *rsbR1*-only (*rsbL* C56A) genotype Δ*rsbR2* Δ*rsbR3* Δ*rsbR4 rsbL* (C56A) and the genotype of *rsbR1*-only (Δ*rsbL*) consists on Δ*rsbR2* Δ*rsbR3* Δ*rsbR4* Δ*rsbL*. Statistical analysis was performed using a paired student *t* test relative to the untreated *rsbR1*-only (*rsbL* C56A) after 15 min, for RT-qPCR data and untreated wild type at time 0 min, for acid challenge data. (*, *P* value of <0.05; **, *P* value of <0.01; ***, *P* value of <0.001).

**S3 Fig.**
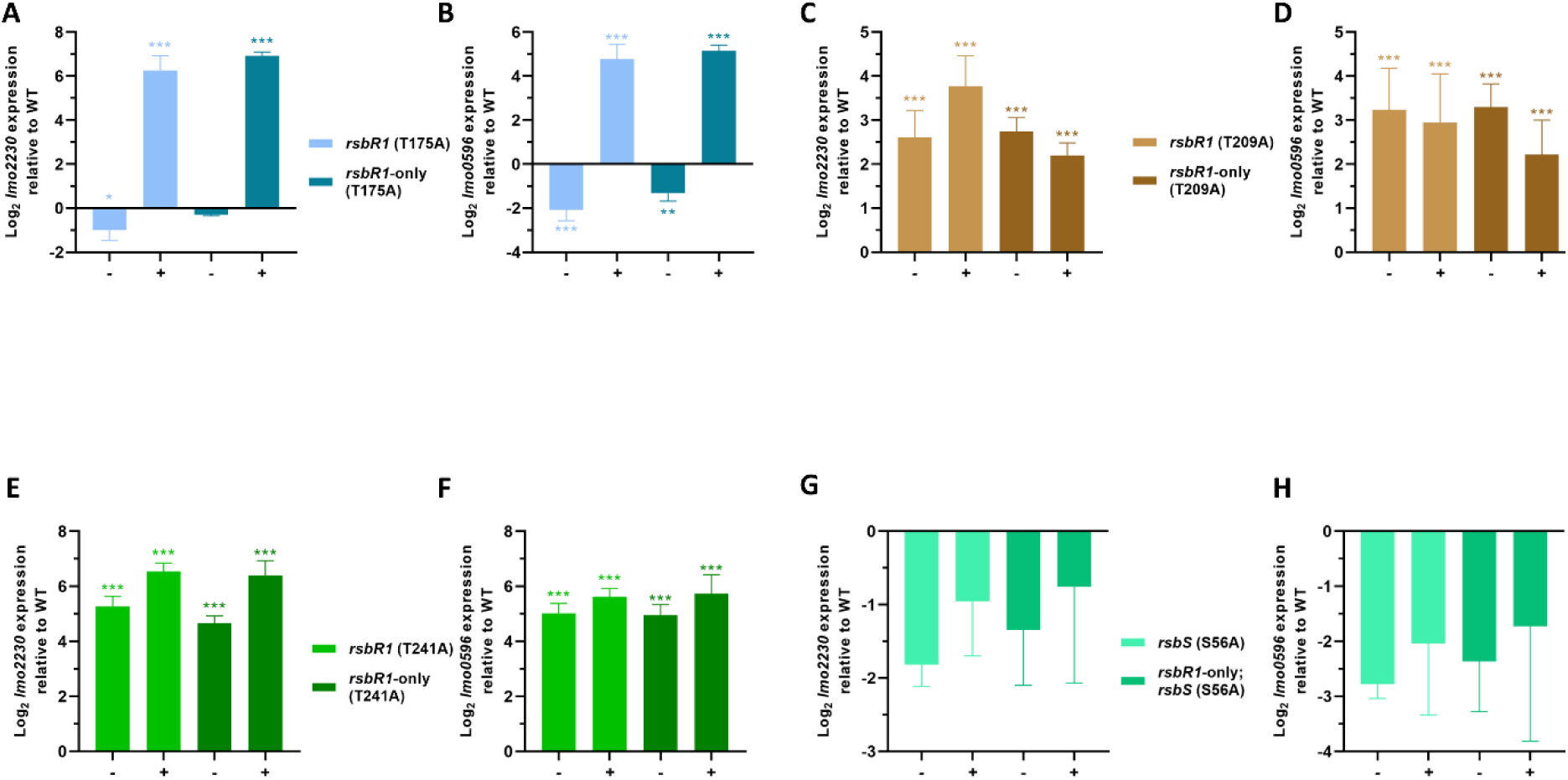
The RsbR paralogues do not interfere with RsbR1 acid sensing capacity. Expression of σ^B^-dependent genes (A, C, E, G) *lmo2230* and (B, D, F, H) *lmo0596*, obtained from mid-log phase cultures of *rsbR1* (T175A), *rsbR1* (T209A), *rsbR1* (T241A), *rsbS* (S56A) and their respective *rsbR1*-only backgrounds. These strains were untreated (-) and treated (+) for 15 min in pH 5.0 at 37°C. The *rsbR1*-only backgrounds consist on the following genotype Δ*rsbR2*; Δ*rsbR3*; Δ*rsbR4*; *rsbL* (C56A). Statistical analysis was performed using a paired student *t* test relative to the untreated wild type strain after 15 min (*, *P* value of <0.05; **, *P* value of <0.01; ***, *P* value of <0.001).

**S4 Fig.**
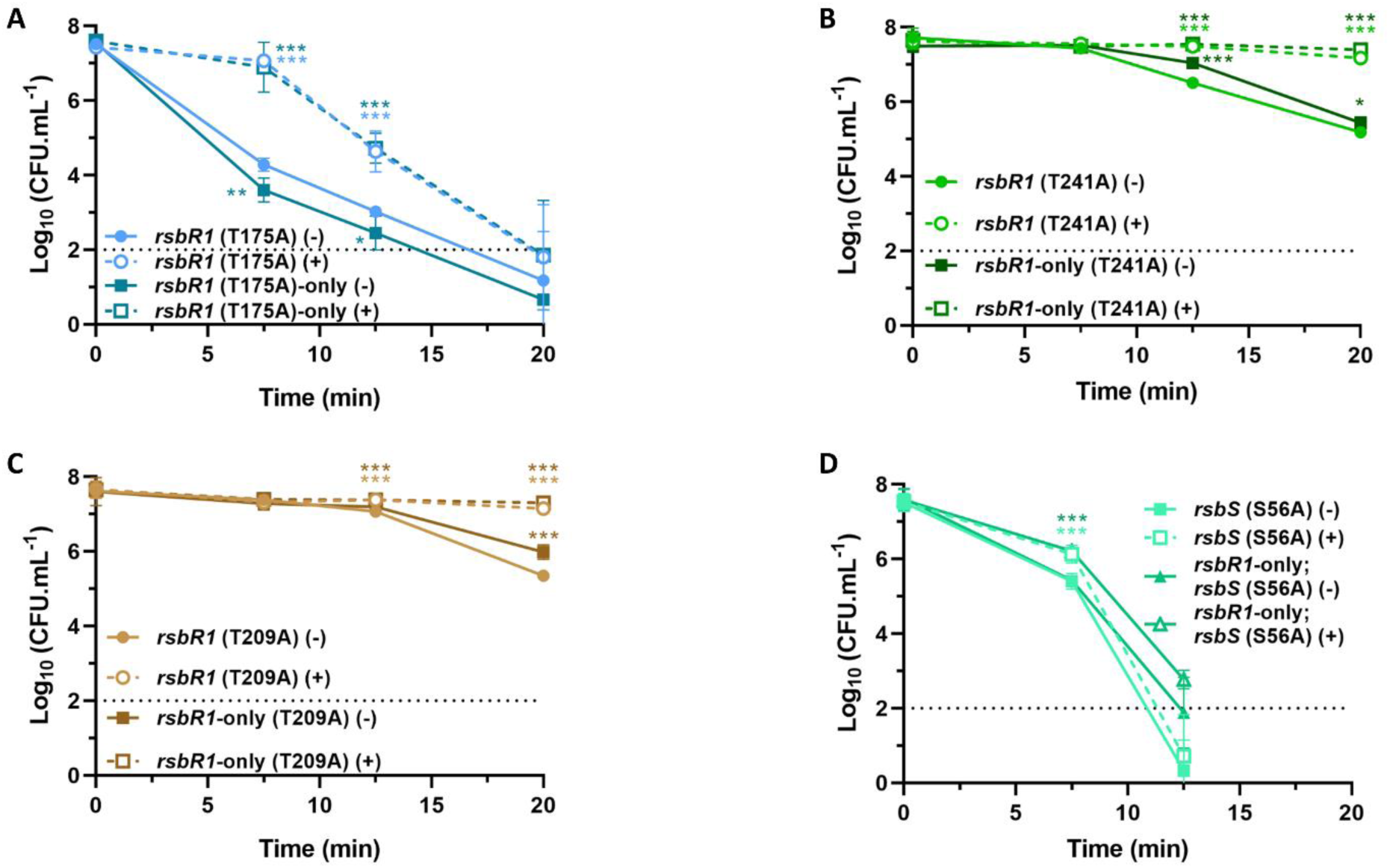
The RsbR paralogues do not affect *L. monocytogenes* acid adaptation. Acid challenge of mid-log phase cultures of *rsbR1* (T175A), *rsbR1* (T209A), *rsbR1* (T241A), *rsbS* (S56A) and their respective *rsbR1*-only backgrounds, treated in BHI acidified to pH 5.0 for 15 min at 37°C. Survival data is expressed as Log_10_ (CFU.mL^-1^). The *rsbR1*-only backgrounds consist in Δ*rsbR2*; Δ*rsbR3*; Δ*rsbR4*; *rsbL* (C56A). The dashed line represents the detection threshold. Samples were taken at 0, 7.5, 12.5 and 20 min. Survival data is expressed as Log_10_ (CFU.mL^-1^). Statistical analysis was performed using a paired student *t* test relative to the respective untreated parental strain after at time 0 min (*, *P* value of <0.05; **, *P* value of <0.01; ***, *P* value of <0.001).

**S5 Fig.**
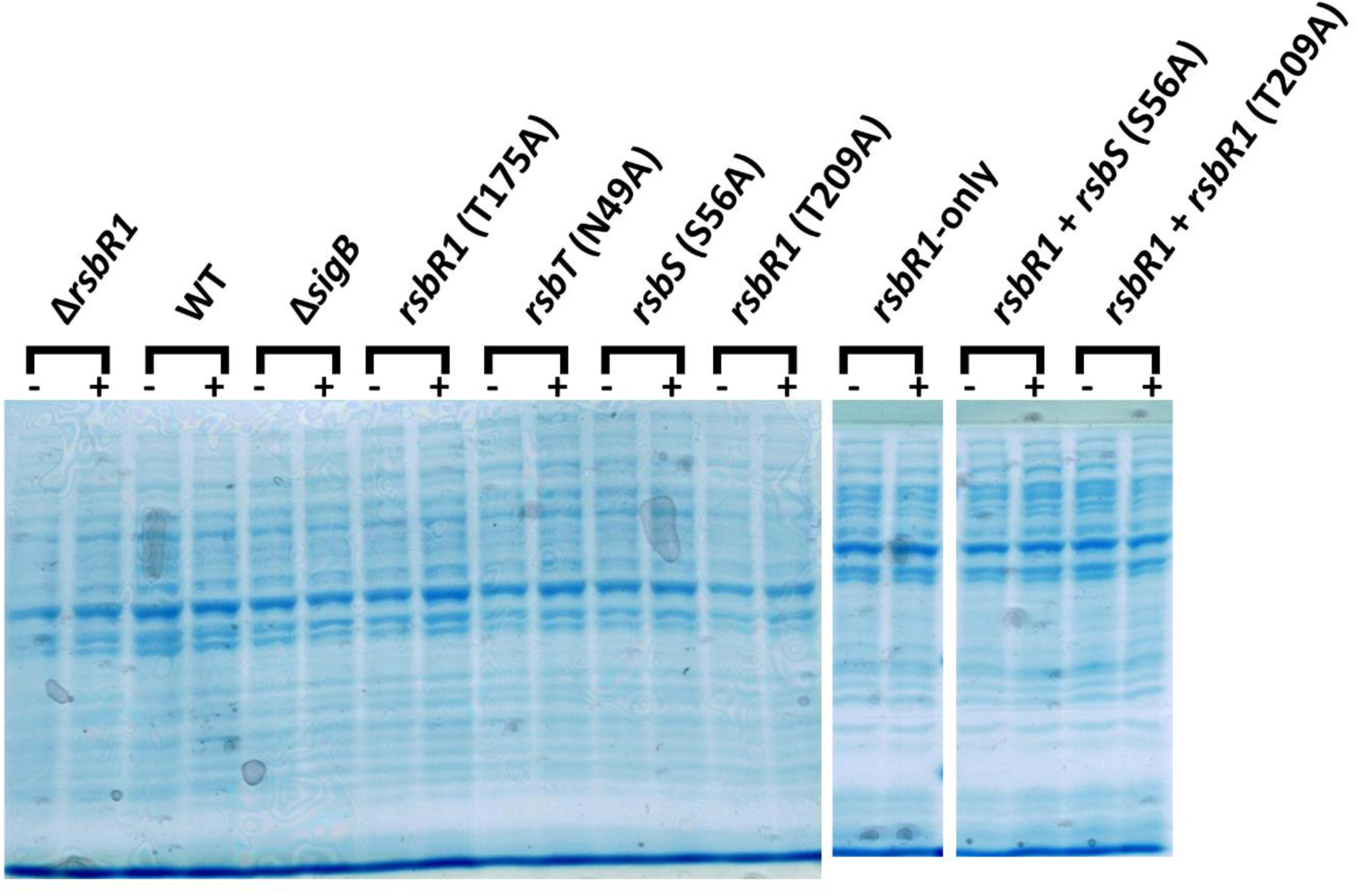
Loading control of samples used for the phosphorylation analysis. SDS-PAGE gels of normalized total protein extracts used for the Phos-tag and western-blots made in this study. Total protein extractions were obtain from mid-log phase cultures untreated (-) and treated (+) in pH 5.0 for 15 min at 37°C. Each protein sample was obtained from ∼7 x 10^7^ cells. Total protein was separated in 12% acrylamide/bis-acrylamide gels and subsequently stained with Coomassie dye.

**S6 Fig.**
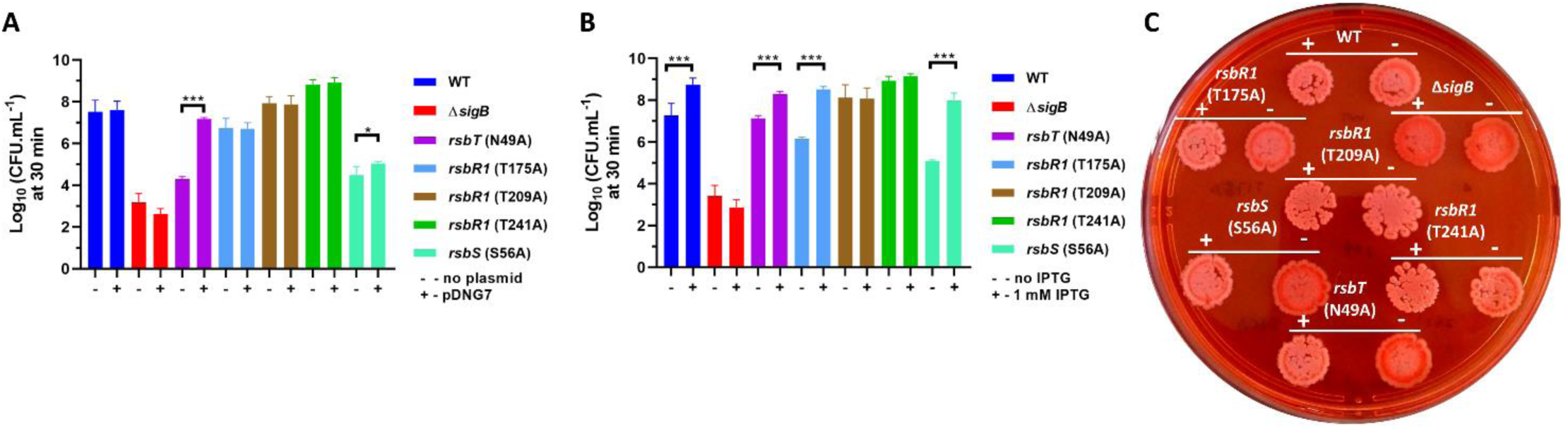
Genetic complementation with *rsbRST* restores signal transduction in stressosome inactive strains. Mutant strains were genetically complemented with pDNG7, which contains the native *rsbR rsbS rsbT* open reading frame under the control of a leaky IPTG inducible promoter (P*_help::lacOid_*). (A) Complemented (+) and non-complemented (-) strains were grown to stationary phase without IPTG in BHI at 37°C and then challenged in BHI acidified to pH 2.5. (B) Complemented strains were grown with (+) and without IPTG (-) and subsequently challenged to pH 2.5. Results are expressed in Log_10_ (CFU.mL^-1^). (C) Stationary phase cultures were plated in BHI agar containing 25 µg.mL^-1^ of Congo Red and 1 mM of IPTG and further incubated at 37°C for 24 h and transferred to 30°C for more 3 days.

